# Alcohol Responsive Genes in the Neutrophils of Acute-On-Chronic Liver Failure Reveal Modulation of Intracellular Calcium, ROS, and Phagocytosis

**DOI:** 10.1101/2023.08.16.553495

**Authors:** Rohini Saha, Ditam Chakraborty, Aditya Roshan, Shalini Sharma, Tamizhini Loganathan, Shalimar, Pragyan Acharya

## Abstract

Acute-on-chronic liver failure (ACLF) is characterized by rapid depletion of hepatic reserves, innate immunity-driven pathogenesis and high short-term mortality. Elevation of neutrophil counts is associated with mortality in alcoholic-ACLF. In India, one of the most common etiologies in ACLF for both chronic and acute injury is alcohol. We used a combination of clinical transcriptomics and a novel *in vitro* laboratory model to look at gene signatures unique in alcoholic ACLF patients and study phenotypic changes upon alcohol treatment. CCL20 and IL1β were two of the top overexpressed genes in alcoholic-ACLF which were validated in the alcohol-neutrophil model, using neutrophils isolated from CLD patients (acute-on-chronic model), and healthy individual (acute model). In the alcoholic-neutrophil models, 300 mg/dL dose of ethanol for 24 hours, caused reduced ROS generation, reduced phagocytic ability and dampening of intracellular calcium reserves. We conclude that the effects of alcohol at least partially explain immune dysfunction in ACLF such as dampening of ROS and phagocytosis in neutrophils while retaining inflammatory roles such as expression of CCL20 and IL1β. Systematic curation of previously published alcohol related liver tissue transcriptomic data also confirmed overlapping genes with alcoholic-ACLF circulating neutrophils. Docking of ethanol with CCL20 and IL1β showed spontaneous binding.

## 1. INTRODUCTION

Acute-on-chronic liver failure (ACLF) results when an acute injury occurring over a chronic liver disease causes rapid depletion of hepatic reserves [1]. The hallmarks of ACLF include systemic inflammation, multiple organ dysfunction and high short-term mortality [2]. The pathogenesis of ACLF is driven by activation of a dysfunctional innate immune response, causing cytotoxicity and systemic inflammation. In recent years, neutrophil to lymphocyte ratio (NLR) has been shown to be an important predictor of outcomes in ACLF [3,4]. In addition, independent groups have reported various molecular defects in the polymorphonuclear cells (neutrophils) derived from ACLF patients [5,7,8,9]. However, the detailed nature and cause of such dysfunction is not yet known. As a result, a systematic preventive or therapeutic strategy to avoid such innate immune dysfunction does not exist. A major challenge in understanding the causes of immune pathogenesis in ACLF, is the heterogeneity of the disease. ACLF is characterized in terms of variable clinical definitions across continents, varied chronic and acute etiologies, presence or absence of sepsis and the involvement of multi-organ dysfunction [7]. While these variables are characteristic of ACLF and cannot be completely avoided, they warrant the design of model systems and experimental strategies that allow us to delineate the effects of individual triggers of the innate immune pathogenesis in ACLF.

Alcohol is a major etiology for ACLF globally, both for the chronic disease component as well as acute injury [10]. Patients suffering from alcohol associated ACLF (alcoholic-ACLF) are more susceptible to bacterial infections due to defects in their innate and adaptive immune responses [4]. Translocation of bacterial LPS from the gut to the circulation upon alcohol consumption and sensitization of liver and extra hepatic organs, is believed to lead to excessive cytokine release and tissue injury in alcoholic-ACLF that can worsen into sepsis leading to mortality. In the present study, we hypothesized that alcohol is one of the triggers for pathogenic neutrophil responses in ACLF. Therefore, our objective was to investigate the influence of alcohol on gene expression and function of neutrophils using a combination of approaches which included re-analysis of published neutrophils transcriptome, validation of select genes in an independent ACLF patient group and the use of a novel *in vitro* model system representing an acute alcoholic condition, and an acute-on-chronic alcoholic condition, to examine the effect on neutrophils phenotypic function.

## 2. MATERIALS AND METHODS

### 2.1 NEUTROPHIL TRANSCRIPTOMICS OF ALCOHOLIC VS NON-ALCOHOLIC ACLF PATIENTS

The neutrophil transcriptome data was retrieved from the National Center for Biotechnology Information’s (NCBI) Gene Expression Omnibus (GEO) database (GSE156382) based on our previously published study [7]. Patient characteristics and sample preparation methods are briefly outlined below-

#### Patient characteristics

Acute-on-chronic liver failure (ACLF) was defined as per the APASL [6] guidelines. Briefly, patients were recruited at the Department of Gastroenterology, AIIMS, New Delhi, India and blood samples were collected with prior informed consent. 12 ACLF patients were randomly chosen for a microarray experiment. In the microarray experiment analysis, ACLF patients were stratified based on alcoholic etiology (Alcoholic ACLF n=5, Non-alcoholic ACLF n=7). The Alcoholic ACLF patients had both the acute and chronic etiology as alcohol related, which included long-term alcohol consumption as well as alcohol binge. Patient baseline characteristics are provided in Table 1.

**Table 1.** Baseline characteristics of ACLF (n=12) patients used in Microarray experiment. The 12 ACLF samples used in microarray experiment GSE156382, were stratified as alcoholic ACLF (n=5) and Non-alcoholic ACLF (n=7). Baseline patient characteristics were compared and due to a small sample size, unpaired T-test was performed for all the variables. Values are mentioned as Mean ± SD (except for male percentage).

|  | Non-Alcoholic ACLF (n=5) | Alcoholic ACLF (n=7) | P value |
| --- | --- | --- | --- |
| Age, years, Mean ± SD | 50 ± 6.0 | 39 ± 8.0 | 0.35 |
| Males % | 100% | 71.40% | 0.20 |
| Hemoglobin (g/dL) | 9.10 ± 1.30 | 9.20 ± 1.10 | 0.94 |
| Sodium (mEq/L) | 140.8 ± 4.5 | 138.0 ± 4.4 | 0.70 |
| Albumin (g/dL) | 2.16 ± 0.30 | 2.70 ± 0.30 | 0.34 |
| Total Leukocyte Count (x10<sup>3</sup> per mm<sup>3</sup>) | 13.75 ± 3.12 | 14.32 ± 3.81 | 0.92 |
| Platelet (x10<sup>3</sup> per mm<sup>3</sup>) | 79.20 ± 35.0 | 67.80 ± 16.40 | 0.74 |
| Urea (mg/dL) | 114.0 ± 51.60 | 98.70 ± 32.30 | 0.79 |
| Creatinine (mg/dL) | 3.60 ± 1.80 | 2.0 ± 0.46 | 0.30 |
| Potassium (mEq/L) | 4.30 ± 0.60 | 4.0 ± 0.50 | 0.70 |
| Bilirubin (mg/dL) | 6.12 ± 1.50 | 20.11 ± 6.20 | 0.13 |
| Aspartate Aminotransferase (IU/L) | 169.0 ± 63.0 | 560.0 ± 446.20 | 0.57 |
| Alanine Transaminase (IU/L) | 102.0 ± 52.50 | 195.5 ± 140.80 | 0.66 |
| Serum Alkaline Phosphatase (IU/L) | 637.70 ± 249.40 | 309.0 ± 75.70 | 0.13 |

#### Sample preparation

Blood samples were processed for neutrophils enrichment, as stated previously [7]. The resulting neutrophils layer was collected, washed and used for mRNA preparation. Microarray sample preparation was done according to the Agilent manufacturer’s protocol. The Agilent SurePrint G3 human gene expression array kit (G4851C) was used and data was analysed with Feature Extraction Software 12.1.0.3 (Agilent) using default parameters. The processed signal intensities were further used for gene expression analysis. The processed metadata is submitted to GEO Database and can be accessed with the number GSE156382 [7].

Validation of gene expression (CCL20 and IL1β) were assessed in ACLF (n=30), CLD (n=10), Healthy control (n=15). The ACLF patient group was further stratified into Alcoholic ACLF and Non-alcoholic ACLF. Baseline characteristics are provided in Supplementary Table S3.

### 2.2 IN VITRO MODEL TO ASSESS EFFECTS OF ALCOHOL ON NEUTROPHIL GENE EXPRESSION AND PHENOTYPE

In order to study the effects of alcohol on neutrophils gene expression and phenotype directly, we developed an *in vitro* model system (Figure 1). Neutrophils were enriched using the double gradient system as stated before [7]. Enriched neutrophils were seeded on a 6 well plate at a density of 1 × 10^5^ cells. Complete RPMI media was supplemented with 300 mg/dL of culture-grade 100% ethanol. The plates were incubated for 24 hours at 5% CO_2_, and 37°C. Post Incubation, the neutrophils were harvested by centrifugation of the collected media and washed twice with incomplete RPMI. The resulting pellet was lysed using TRIzol reagent, and proceeded to RNA isolation. The total RNA from neutrophils before and after alcohol incubation were estimated using NanoDrop (Thermo Scientific) and 500 ng RNA was converted to cDNA using the Thermo Scientific Verso kit (AB-1453 A). The primers for the genes *CCL20* and *IL1β* were optimized and qRT-PCR was performed using the pre- and post-alcohol incubation cDNA. Data was represented as log Fold change. The data reported here are according to MiQE guidelines (refer MiQE docx in supplementary files) (Bustin et al. 2009).

**Figure 1.**
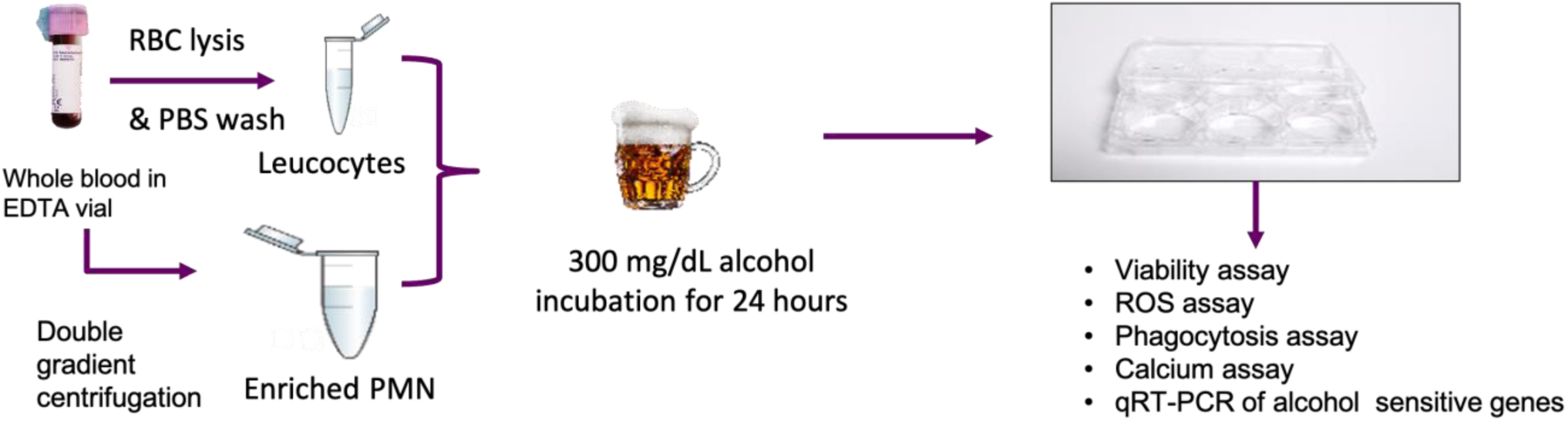
Methodology outline of *in vitro* alcohol treated neutrophil model. A dose of 300 mg/dL for 24-hour duration was used for all assays. Whole blood was processed for neutrophil enrichment by double gradient centrifugation. Neutrophils were used for qPCR analysis. Whole blood was directly used for neutrophil phenotypic assays. Blood sample from healthy individuals was used as a representation of acute alcohol model and blood sample from CLD individuals was used as a representation of acute-on-chronic model.

For phenotypic assays, 100ul whole blood was incubated with RPMI media supplemented with alcohol for 24 hours at 5% CO_2_ and 37°C. Post alcohol incubation, cells were harvested with a brief trypsinization. Harvested total blood cells were treated with RBC lysis buffer and the leucocyte pellet was further used for flow cytometric analysis. Intracellular calcium reserve was measured using Fluo-4 Direct™ Calcium assay reagent (F10471, Invitrogen) as per the manufacturer’s protocol. Phagocytosis capacity was assessed by AF488 conjugated *E. coli* particles (E13231, Thermo Kit). For ROS assay, the dye DHR123 (D1054, Sigma-Aldrich) was directly added onto the 6 well plate containing the whole blood cells, before trypsinization followed by incubation for 30 mins in dark, at 5% CO_2_, and 37°C. The cells were washed and reconstituted in FACS buffer and proceeded with flow cytometry acquisition. Data was analysed using FlowJo™ software [21].

### 2.3 IN SILICO INTERACTION STUDY OF ETHANOL WITH THE PROTEINS OF INTEREST

Further, in order to understand the nature of interactions of ethanol with cell surface receptors (GPCRs), cytokines and intracellular transcription factors, molecular dockings were performed. CCL20 and IL1β were selected as receptors with ethanol as the ligand. Three-dimensional structures of the receptors proteins and ligand were downloaded from RCSB Protein Data Bank [22] and PubChem [23], respectively. The structure files (in “.pdb” format) of the receptors were opened in PyMol 2.5.2 [24] for removal of the bound ligands and water molecules.

Molecular docking was performed using Autodock 4.2 software [25]. At first, the receptor proteins were prepared for docking by adding Kolmann charges, repairing missing atoms (if any) and addition of polar hydrogen atoms. For the ligand ethanol, energy minimization was performed by adding Gasteiger charges and minimum number of torsions was set to 1. Since ethanol has been known to interact non-specifically to many protein grooves, blind docking was performed by setting the grid box dimensions to the maximum values so that it covered the entire receptor. Blind docking algorithm enables the ligand to scour for compatible binding sites on the receptor surface and bind with the site having the highest degree of complementarity [26]. The docking parameters were set to Genetic Algorithm with the number of runs as 20 and a population size of 300. The resulting receptor-ligand docked complexes were saved in the “.pdb” format and analyzed using BIOVIA Discovery Studio 2021 software [27].

### 2.4 INTEGRATIVE CURATION OF LIVER TISSUE TRANSCRIPTOMICS OF ALCOHOLIC LIVER DISEASE

Human gene expression of Alcohol liver disease datasets was retrieved from the National Center for Biotechnology Information’s (NCBI) Gene Expression Omnibus (GEO) database (http://www.ncbi.nlm.nih.gov/geo/), Arrayexpress (https://www.ebi.ac.uk/arrayexpress/), and OmicsDI (https://www.omicsdi.org/), by searching the keywords “Alcoholic liver disease” and “Alcoholic Hepatitis” from organism “*Homo sapiens*”. The experiment types used in this study were “Expression profiling by array” and “Expression profiling by High-throughput Sequencing”. Gene expression datasets were filtered based on inclusion and exclusion criteria.

#### Inclusion criteria

i) The gene expression profile was derived from human liver tissue of Alcoholic liver disease from the experiment of Expression profile by array and Expression profile by high throughput sequencing, (ii) Sample datasets were to have both cases and control groups, (iii) The cases group should contain the Alcohol liver disease patients and control group should have healthy subjects, (iv) Each case and control groups were to have at least three samples.

#### Exclusion criteria

i) No HCC samples or other cancer related datasets were selected, (ii) non-liver tissue datasets were excluded, (iii) Organisms other than *Homo sapiens* were excluded.

Based on the above conditions, the five datasets were chosen for further data analysis.

Among the shortlisted five datasets, GSE159676 [11] and GSE10356 [12] were further excluded. The dataset GSE159676 had only one alcohol liver disease sample and 6 normal samples. The dataset GSE10356 had a customized Liverpool platform of transcriptomic experiment, which could not be merged with the data of other platforms. Hence a total of two datasets were selected for further analysis from the above criteria. The selected datasets were GSE155907 [13] and GSE28619 [14]. From these datasets GSE155907 was Expression Profiling by High throughput Sequencing (RNA-Seq) and GSE28619 was Expression profiling by microarray. The dataset of GSE155907 had a total of 9 samples (4 control groups and 5 Alcohol-related groups and the dataset of GSE28619 had 22 Samples (7 control groups and 15 Alcoholic-related groups).

### 2.5 OVERLAPPING GENE EXPRESSION SIGNATURES ACROSS CIRCULATING NEUTROPHILS IN ACLF AND LIVER TISSUE OF ALCOHOLIC LIVER DISEASE

The differential expressed gene list from neutrophils microarray data was analyzed using DAVID [16] for gene ontology classification and Metascape [17] for pathway enrichment analysis. The differentially expressed genes (DEG) from the two published dataset was analyzed using WebGestalt [18]. We further looked at the overlapping genes between our experimental neutrophil datasets, and the DEGs of the two published datasets, GSE28619, GSE155907 and alcohol liver disease related genes-published on the database DisGeNET [19]. It is a database for classification of genes based on their association to any particular disease. We extracted the gene set associated with alcoholic liver disease and compared with our neutrophils microarray data to find a common gene signature influenced by alcohol. The overlapping genes are represented by Venn diagrams using online tool Venny 2.1.0 [20].

## 3. RESULTS

### 3.1 DIFFERENTIAL GENE REGULATION AND SIGNALING PATHWAY IN ALCOHOLIC ACLF VS NON-ALCOHOLIC ACLF PATIENT DERIVED NEUTROPHILS

Our previously published neutrophils transcriptomic dataset GSE156382 [7] was reanalysed to look at gene expression profiles altered in neutrophils due to alcohol. Briefly, the ACLF patient cohort (n=12) was stratified as Alcoholic-ACLF and Non-Alcoholic ACLF (including Viral hepatitis, autoimmune hepatitis, cryptogenic causes, etc.). The alcoholic ACLF group had 2 sepsis and 3 sterile inflammation cases and the non-alcoholic ACLF group had 4 sepsis and 3 sterile inflammation cases. The baseline patient characteristics are presented in Table 1.

The microarray signal intensity output files, FEF (feature extraction files for Agilent) were submitted to the GEO analysis software and samples were assigned the tag ‘Test’ (Alcoholic ACLF) and ‘Control’ (Non-alcoholic ACLF). The results were exported in .XLS format and sorted based on the Fold change and p-value of significance. After setting the cut-off of ≤ 2 ≥ for Fold change, and p-value ≤ 0.05, a total of 283 genes were found to be differentially expressed in neutrophils of Alcoholic ACLF and Non-alcoholic ACLF patients, with 223 genes being downregulated and 60 genes being upregulated (Figure 2, A-B), (Supplementary Table S1). The top enriched pathways were T-cell differentiation (Log p-value= −6.47), GPCR signaling (Log p-value= −5.27), Calcium Signaling (Log p-value= −3.29), Neurotransmitter release (Log p-value= −2.71), transmembrane receptor tyrosine kinase (Log p-value= −5.21) and cytokine signaling (Log p-value= −4.04). The pathways with the gene entity symbols and regulations are represented in Figure 2D. (Supplementary Table S2)

**Figure 2.**
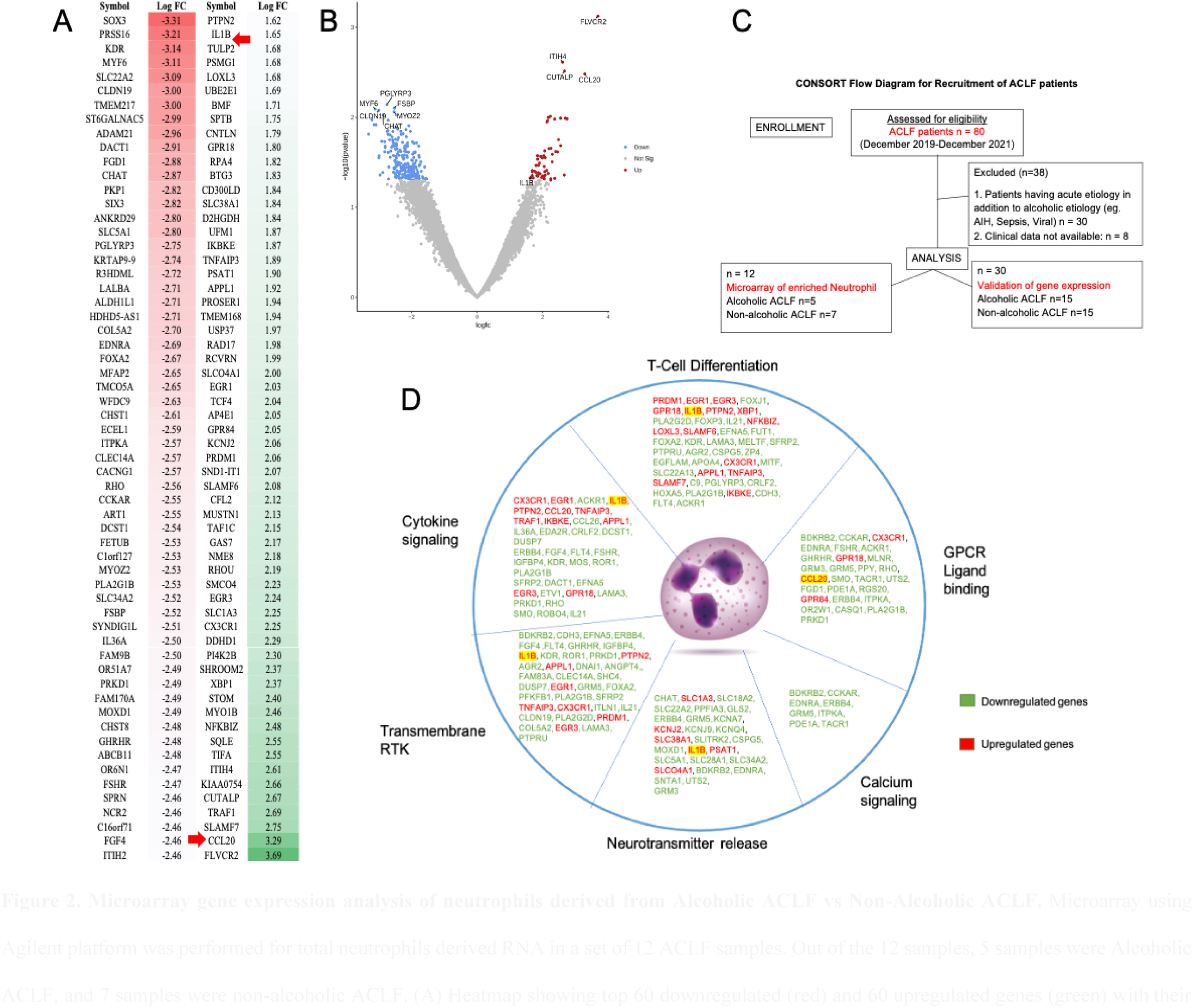
Microarray gene expression analysis of neutrophils derived from Alcoholic ACLF vs Non-Alcoholic ACLF. Microarray using Agilent platform was performed for total neutrophils derived RNA in a set of 12 ACLF samples. Out of the 12 samples, 5 samples were Alcoholic ACLF, and 7 samples were non-alcoholic ACLF. (A) Heatmap showing top 60 downregulated (red) and 60 upregulated genes (green) with their respective log Fold change values, (B) Volcano plot of LogFC vs –Log(p-value) depicting the differentially expressed genes. The downregulated genes are represented as blue dots and upregulated genes are represented as red dots, (C) CONSORT flow diagram showing the recruitment of ACLF patients for microarray experiment and validation experiment, (D) Gene Ontology analysis of DE genes of neutrophils isolated from Alcoholic-ACLF Patients. Summarizing the results of Gene Ontology (GO) analysis of the 283 DE genes of neutrophils isolated from ACLF patients with alcohol etiology. The online tool DAVID was used for performing GO analysis. Threshold count was set to 2 and EASE score was set to 0.1.

### 3.2 qRT-PCR VALIDATION OF CCL20 AND IL1β GENES IN IN VITRO ALCOHOL NEUTROPHILS MODEL AND ACLF PATIENT DERIVED NEUTROPHILS

Enriched neutrophils from ACLF patients, CLD patients, as well as healthy blood samples were used for qPCR assays. After enrichment, neutrophil population was found to be 85.45% (± 6.20%), with 1.14% (± 1.02%) eosinophil/basophil and 1.9% (± 2.00%) NK cell contamination (Supplementary Figure S1). In addition, neutrophils viability assay results of the enriched neutrophils showed that on an average 91.25% cells (% of total neutrophils) were viable before incubation with alcohol (Supplementary Figure S2, top panel) and on an average 95.67% of cells (% of total neutrophils) were viable after incubation with alcohol (Supplementary Figure S2, bottom panel).

The neutrophil *in vitro* acute alcoholic model was carried out using enriched neutrophils from healthy individuals (n=14) and CLD (n=14). Both *IL1β* and *CCL20* from enriched healthy neutrophils (acute model) and CLD neutrophils (acute-on-chronic model) after 24 hours of incubation with alcohol showed significant upregulation compared to pre-incubation samples. Mean (**±** SD) gene expression of IL1β for alcoholic neutrophils model pre-incubation sample group was 0.24 (**±** 2.70) and for post alcohol sample group was 11.09 (**±**11.64); p-value = 0.0022 (Figure 3A). Mean (**±** SD) gene expression of CCL20 for alcoholic neutrophils model pre-incubation sample group was −0.21 (**±** 5.10) and for post alcohol sample group was 12.28 (**±** 11.24); p-value < 0.0008 (Figure 3B). In the in vitro acute-on-chronic alcohol model, using CLD samples, however, only CCL20 was significantly upregulated post alcohol incubation, with a mean (± S.D.) of gene expression of pre incubation: 0.35 (± 2.82) and post incubation: 3.37 (± 2.38); p-value = 0.005 (Figure 3D). Alcohol did not induce *IL1β* upregulation in CLD neutrophils: mean (± SD) gene expression of pre incubation: −0.09 (± 3.11) and post incubation: 0.68 (± 4.4); p-value = 0.60 (Figure 3C). In order to validate the microarray results, qPCR experiments were performed to compare expressions of IL1β and CCL20 among ACLF, CLD, and healthy controls (Figure 3 E-F) and alcoholic vs non-alcoholic ACLF patients (Figure 3 G-H). IL1β expression in neutrophils was significantly different in CLD vs Healthy [Mean (± S.D): 0.52 (± 1.66) vs −8.19 (± 2.63); p-value < 0.0001], and ACLF vs Healthy [Mean (±S.D.): 2.79 (± 0.74) vs −8.19 (± 2.63); p-value < 0.0001] (Figure 3E). Whereas, CCL20 expression did not show any significant alteration among, ACLF patients [−0.09 (± 3.44], CLD [(2.44 (± 4.82)] and healthy [−0.01 (± 5.09)]; ANOVA p-value =0.24 (Figure 3F). The mean (± SD) gene expression of IL1β for Non-alcoholic ACLF was found to be −0.004 (± 3.07) while for Alcoholic ACLF it was −0.81 (± 4.93); p-value = 0.59 (Figure 3G). The mean (± SD) gene expression of CCL20 for Non-alcoholic ACLF was −0.003 (± 2.42) while for Alcoholic ACLF was −0.13 (± 3.88); p-value = 0.91 (Figure 3H). This finding did not corroborate with the microarray results, which might be due to patient heterogeneity within the defined alcoholic and non-alcoholic groups. The baseline characteristics for the alcoholic and non-alcoholic ACLF patients have been summarized in Supplementary Table S3.

**Figure 3.**
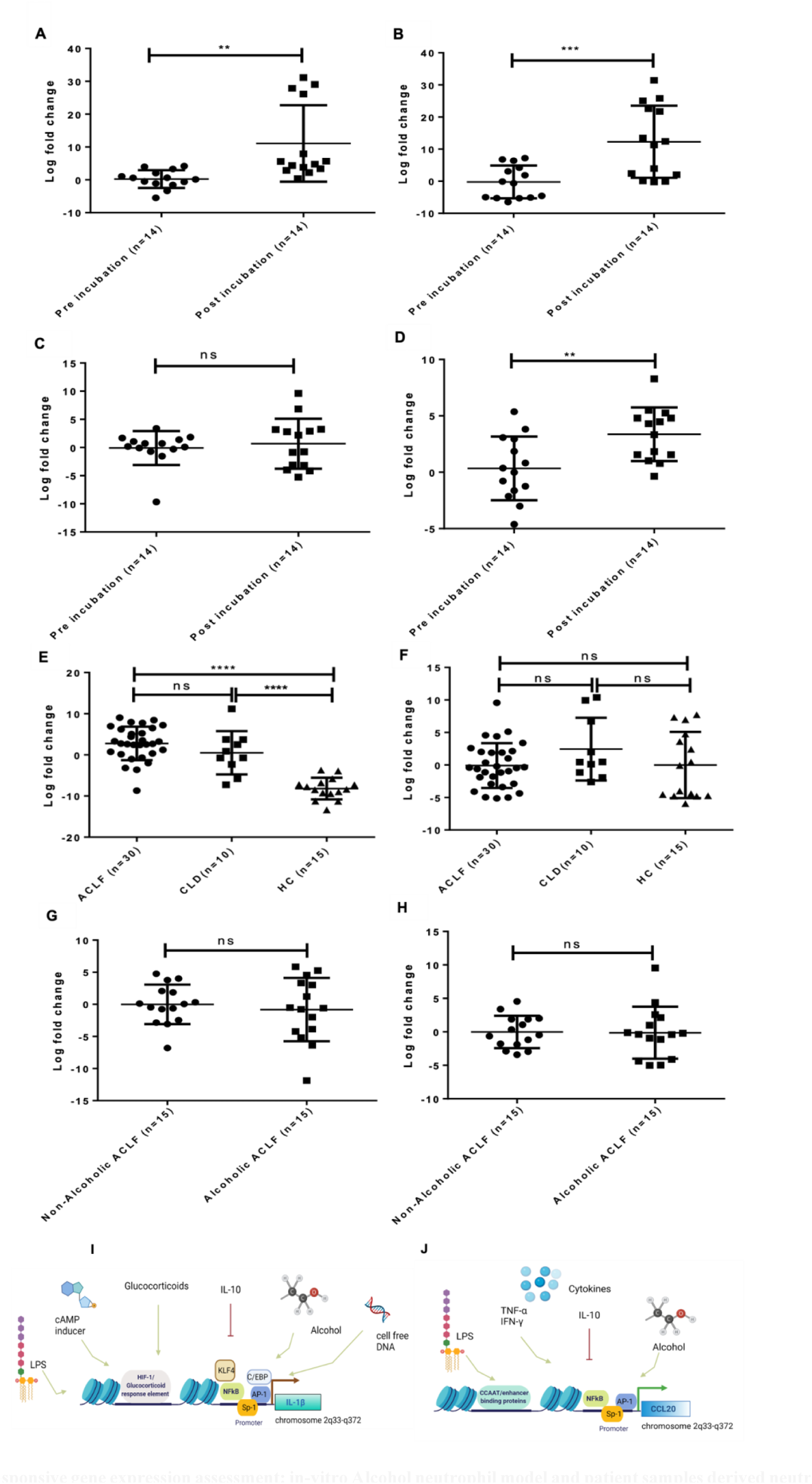
Alcohol responsive gene expression assessment: in-vitro Alcohol neutrophil model and patient samples derived neutrophil. qPCR validation of IL1β and CCL20 in the *in vitro alcohol* model (A-B) using healthy neutrophils (acute alcoholic model), (C-D) using CLD neutrophils (acute on chronic alcoholic model). (A) Mean (± SD) gene expression (delta C_T_) of IL1β for acute alcoholic-neutrophils model was 0.24 (**±** 2.70) pre-incubation with EtOH vs 11.09 (**±**11.64) post-incubation with EtOH; p-value < 0.0022 (n=14 for each group). (B) Mean (± SD) gene expression (delta C_T_) of CCL20 for acute alcoholic-neutrophils model was −0.21 (**±** 5.10) pre-incubation with EtOH vs 12.28 (**±** 11.24) post-incubation with EtOH; p-value < 0.0008 (n=14 for each group). (C) Mean (± SD) gene expression (delta C_T_) of IL1β for acute-on-chronic alcoholic-neutrophils model was −0.09 (**±** 3.11) pre-incubation with EtOH vs 0.68 (**±** 4.4) post-incubation with EtOH; p-value= 0.60. (D) Mean (± SD) gene expression (delta C_T_) of CCL20 for acute-on-chronic alcoholic-neutrophils model was 0.35 (**±** 2.82) pre-incubation with EtOH vs 3.37 (± 2.38) post-incubation with EtOH; p-value = 0.005 (n=14 for each group). (E) Mean (± SD) gene expression (delta C_T_) of IL1β in patient samples showed significant difference in CLD vs Healthy [Mean (± S.D): 0.52 (± 1.66) vs −8.19 (± 2.63)]; p-value < 0.0001, and ACLF vs Healthy [Mean (±S.D.): 2.79 (± 0.74) vs −8.19 (± 2.63)]; p-value <0.0001. (F) Mean (± SD) gene expression (delta C_T_) of CCL20 in patient samples showed ACLF patients −0.09 (± 3.44), CLD 2.44 (± 4.82) and healthy −0.01 (± 5.09); ANOVA p-value =0.24. (G) Mean (± SD) gene expression (delta C_T_) of IL1β for non-alcoholic ACLF was −0.004 (± 3.07) vs Alcoholic ACLF it was −0.81 (± 4.93); p-value = 0.59 (n=15 for each group). (H) Mean (± SD) gene expression (delta C_T_) of *CCL20* for non-alcoholic ACLF was −0.003 (± 2.42) vs for Alcoholic ACLF was −0.13 (± 3.88); p-value = 0.91 (n=15 for each group). (I) and (J) depict the multi-factorial modulation of *CCL20* and *IL1β* gene expression. Our data shows that alcohol is one of the factors modulating *CCL20* and *IL1β* gene expression. Log fold changes were normalized and compared using 18S as a housekeeping gene in all qPCR experiments.

### 3.3 IN VITRO MODEL FOR EVALUATING THE EFFECT OF ALCOHOL ON CALCIUM RESERVES, ROS AND PHAGOCYTOSIS IN NEUTROPHILS

In order to study how alcohol affects neutrophil phenotype, we acquired blood from healthy donors and CLD patients, and carried out ROS, Phagocytosis and Intracellular calcium assay, before and after incubating with ethanol (300 mg/dL, 24 hours, 37°C and 5% CO_2_). The dosage of alcohol was determined on the basis of reported alcohol dose which leads to the effects of severe intoxication. While this is not a *bona-fide* model of ACLF, this model allows us to study the effects of acute alcohol injury on healthy cells (acute model), and chronic liver disease cells (acute-on-chronic model) and ascertain whether some of the observations that have been reported as part of the innate immune dysfunction in ACLF, can be explained by the effects of alcohol.

Red blood cell lysis was carried out and the resulting WBC pellet was resuspended in 1X PBS and this single cell suspension was used for the phenotype assays. During flow cytometry acquisition, neutrophils were gated based on their FSC-SSC profile as previously reported in literature [34]. Calcium assay was performed using the Fluo4 calcium binding dye, which corresponds to FITC spectral range. Calcium reserve measurements were done at baseline and post 30-minutes incubation with the dye, in dark, at 37°C and 5% CO_2._ The calcium reserve was calculated as: MFI at 30 minutes –MFI at 0 minutes. The pre-incubation healthy samples showed a median MFI (Inter-Quartile Range) distribution of 54070 (44404-60837) and post-alcohol incubation healthy samples showed a median (Inter-Quartile Range) distribution of 19616 (11485-22587) (Figure 4 B,C). The p-value was computed to be 0.028 according to Mann-Whitney test. The pre-incubation CLD samples showed a median (Inter-Quartile Range) distribution of 51644 (41679-72286) and post-alcohol incubation CLD samples showed a median (Inter-Quartile Range) distribution of 12059 (10356-13594) (Figure 4 D,E). The p-value was computed to be 0.028 according to Mann-Whitney test. Hence, alcohol treatment causes dampening of intracellular calcium reserve in both healthy (acute model) and CLD (acute-on-chronic model) neutrophils. Since calcium reserves are central to neutrophil phenotypic activities such as ROS production and phagocytosis, dysregulated calcium reserve is expected to hamper the downstream processes.

**Figure 4.**
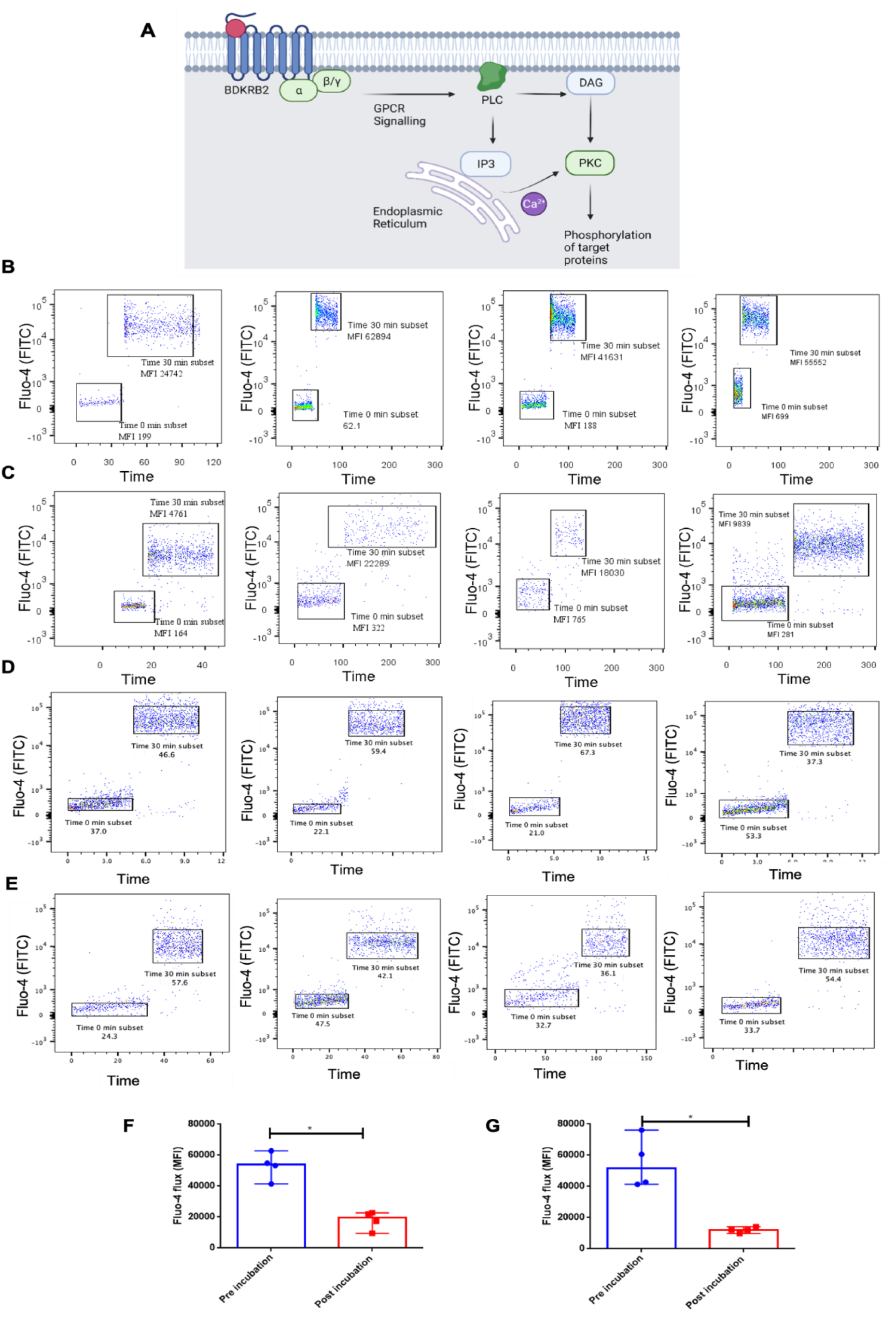
Calcium signaling is dampened in response to alcohol treatment. (A) Several components of the calcium signaling pathway such as GPCRs (BDKRB2, CCKAR, EDNRA, GRM5, TACR1), PDE1A and ITPKA were downregulated in alcoholic-ACLF vs non-alcoholic ACLF suggesting that calcium signaling as a whole might be affected by alcohol. (B-E) Representative images for the measurement of intracellular calcium reserve (MFI) using Fluo4 dye in alcoholic-neutrophils model. Fluo4 dye signal was measured in the FITC channel and represented as FITC-A vs Time plot, indicating the MFI at 0 min and 30 minutes. Intracellular calcium reserve was calculated as MFI at 30 min - MFI at 0 min [Median (Inter-Quartile Range)] (B) Pre-incubation healthy neutrophils (-EtOH): 54070 (44404-60837) (C) Post-alcohol incubation healthy neutrophils (+EtOH): 19616 (11485-22587) (D) Pre-incubation CLD neutrophils (-EtOH): 51644 (41679-72286) (E) Post-alcohol incubation CLD neutrophils (+EtOH): 12059 (10356-13594) (F) Graphical representation of intracellular calcium reserve in acute alcoholic neutrophil model, as indicated by difference in FITC-MFI at time 30 min - time 0 min. Mann-Whitney test has been applied here. [p value= 0.03] (n=4 per group). (G) Graphical representation of intracellular calcium reserve in acute-on-chronic alcoholic neutrophil model, as indicated by difference in FITC-MFI at time 30 min - time 0 min. Mann-Whitney test has been applied here. [p value= 0.03] (n=4 per group).

ROS assay was performed with healthy and CLD pre- and post-alcohol incubation samples using the DHR123 dye, with or without PMA stimulation. PMA is a potent activator of NF-κB in vitro and works at nanomolar concentration range to induce ROS production in cells. The MFI and percentage of cells positive for DHR123 was considered as ROS producing cells. The MFI of ROS detecting dye is directly proportional to the amount of oxygen species released, whereas the percentage of cells generating the dye reflects the activation status of the cells. In the acute-alcoholic neutrophil model, Median (Interquartile range) for pre-incubation FITC MFI was 2573 (1607-3307) and that for post-alcohol incubation was 1280 (1175-1689); Median (Inter-Quartile Range) for pre-incubation FITC MFI (for DHR123+PMA) was 33690 (33391-42712) and for post-alcohol incubation was 4130 (3135-12145) (Figure 5 B). In the acute-on-chronic alcoholic neutrophil model, Median (Interquartile range) for pre-incubation FITC MFI was 1746 (1637-2012) and that for post-alcohol incubation was 1693 (1542-1874); Median (Inter-Quartile Range) for pre-incubation FITC MFI (for DHR123+PMA) was 3222 (2571-3775) and for post-alcohol incubation was 3992 (2334-4120) (Figure 5 D).

**Figure 5.**
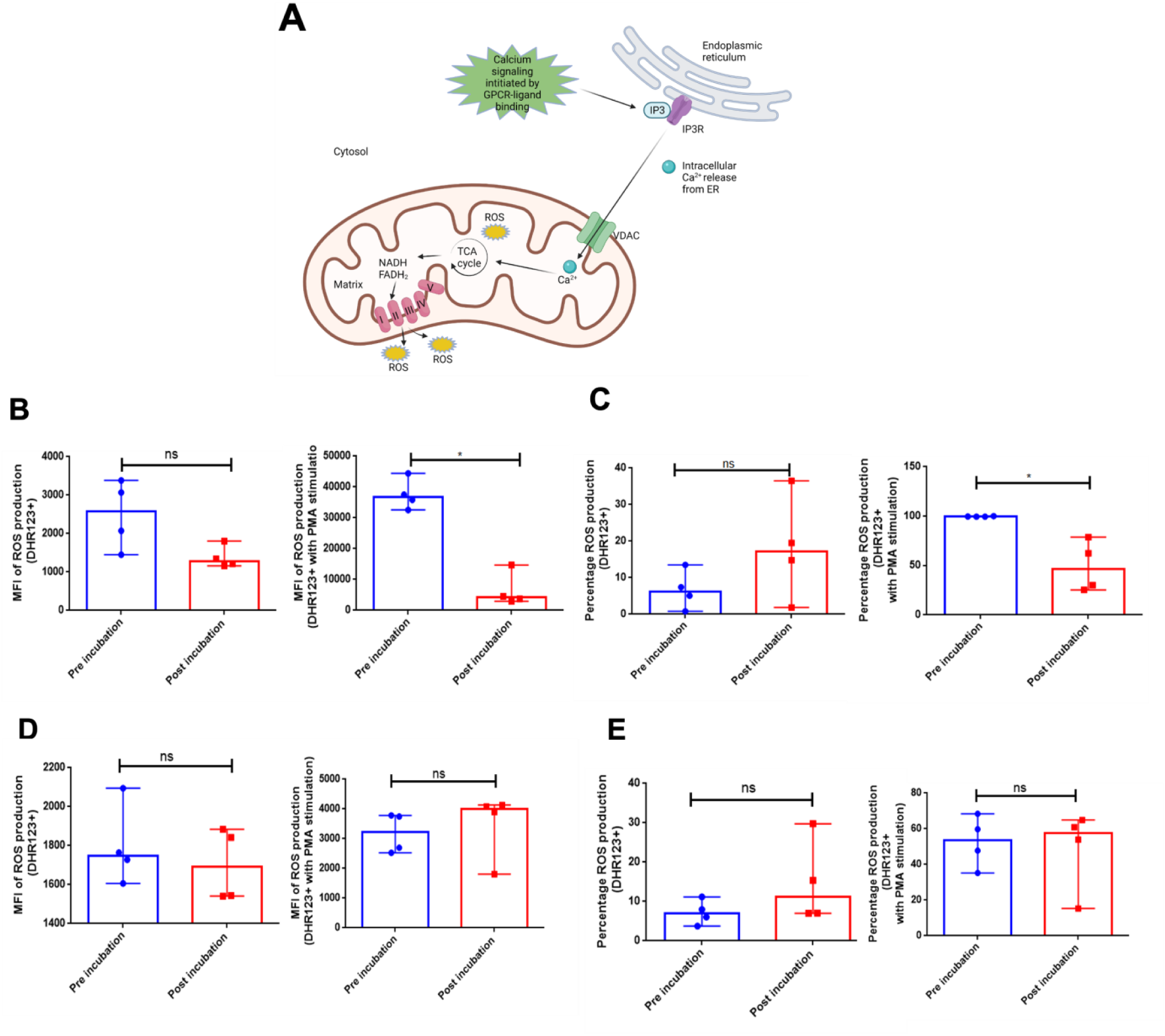
Effect of Alcohol on ROS production. (A) Calcium reserves regulate mitochondrial ROS generation in neutrophils. (B, D) Representative images showing MFI of ROS production using DHR123 dye, with or without PMA stimulation in acute and acute-on-chronic model respectively, (C, E) Representative images showing percentage of cells showing ROS production using DHR123 dye, with or without PMA stimulation acute and acute-on-chronic model respectively. (B) In acute alcohol model, (left) Median (Inter-Quartile Range) of DHR123-MFI in pre incubation vs post alcohol incubation is 2573 (1607-3307) vs 1280 (1175-1689); (right) Median (interquartile range) of DHR123 with PMA MFI in pre incubation vs post alcohol incubation is 36690 (33391-42712) vs 4130 (3135-12145). (C) In acute alcohol model, (left) Median (Inter-Quartile Range) of percentage of cells positive for DHR123 in pre incubation vs post alcohol incubation is 6.24 (1.87-11.98)% vs 17.15 (5.08-32.25)%; (right) Median (Inter-Quartile Range) of percentage of cells positive for DHR123 with PMA in pre incubation vs post alcohol incubation is 99.65 (99.60-99.93)% vs 46.35 (26.53-74.73)%. (D) In acute-on-chronic alcohol model, (left) Median (interquartile range) of DHR123-MFI in pre incubation vs post alcohol incubation is 1746 (1637-2012) vs 1693 (1542-1874); (right) Median (Inter-Quartile Range) of DHR123 with PMA MFI in pre incubation vs post alcohol incubation is 3222 (2571-3775) vs 3992 (2334-4120). (E) In acute-on-chronic alcohol model, (left) Median (Inter-Quartile Range) of percentage of cells positive for DHR123 in pre incubation vs post alcohol incubation is 7.03(4.30-10.39)% vs 11.25 (7.03-26.20)%; (right) Median (Inter-Quartile Range) of percentage of cells positive for DHR123 with PMA in pre incubation vs post alcohol incubation is 53.80 (38.35-66.25)% vs 57.40 (25.03-63.90)%.

There was a significant decrease in intensity of ROS production in healthy neutrophils stimulated with PMA post-alcohol incubation. Thus, alcohol might be involved in hampering the ability of healthy neutrophils to respond to the stimulus of PMA. Whereas, when we took CLD neutrophils, the intensity of ROS production upon PMA stimulation was almost 10-fold lower than that of healthy neutrophils. This dysfunctional phenotype of CLD neutrophils remained unaffected by alcohol treatment.

We also estimated the percentage of cell positives for DHR123, with and without PMA stimulation. In the acute-alcoholic neutrophil model, Median (Inter-Quartile Range) for pre-incubation FITC positive cells was 6.24 (1.87-11.98)% and that for post-alcohol incubation was 17.15 (5.08-32.25)%; Median (Inter-Quartile Range) for pre-incubation FITC MFI (for DHR123+PMA) was 99.65 (99.60-99.93)% and for post-alcohol incubation was 46.35 (26.53-74.73)% (Figure 5C). In the acute-on-chronic alcoholic neutrophil model, Median (Inter-Quartile Range) for pre-incubation FITC MFI was 7.03(4.30-10.39)% and that for post-alcohol incubation was 11.25 (7.03-26.20)%; Median (25%-75%) for pre-incubation FITC MFI (for DHR123+PMA) was 53.80 (38.35-66.25)% and for post-alcohol incubation was 57.40 (25.03-63.90)% (Figure 5 E). It is evident from this finding that CLD neutrophils respond lesser to PMA stimulation as compared to healthy neutrophils, and alcohol does not influence ROS production significantly.

For phagocytosis, the percentage of cells staining for the AF488 marker conjugated to *E. coli* bioparticles (E13231), were considered as phagocytic cells. The percentage of phagocytic cells pre- and post-alcohol incubation were plotted and significance was computed. In the acute alcoholic neutrophils group Median (Inter-Quartile Range) for pre-incubation sample group was calculated as 47.9 (36.0-63.1)%, and for the alcohol incubation sample group as 17.7 (14.78-19.52)% (Figure 6 B). In the acute-on-chronic alcoholic neutrophils group Median (Inter-Quartile Range) for pre-incubation sample group was calculated as 42.3 (41.8-52.10)%, and for the alcohol incubation sample group as 34.55 (25.05-35.73)% (Figure 6 C). Thus, alcohol causes reduction of phagocytosis in both healthy and CLD neutrophils. The extent of reduction is more in healthy (acute model) as compared to CLD (acute-on-chronic model).

**Figure 6.**
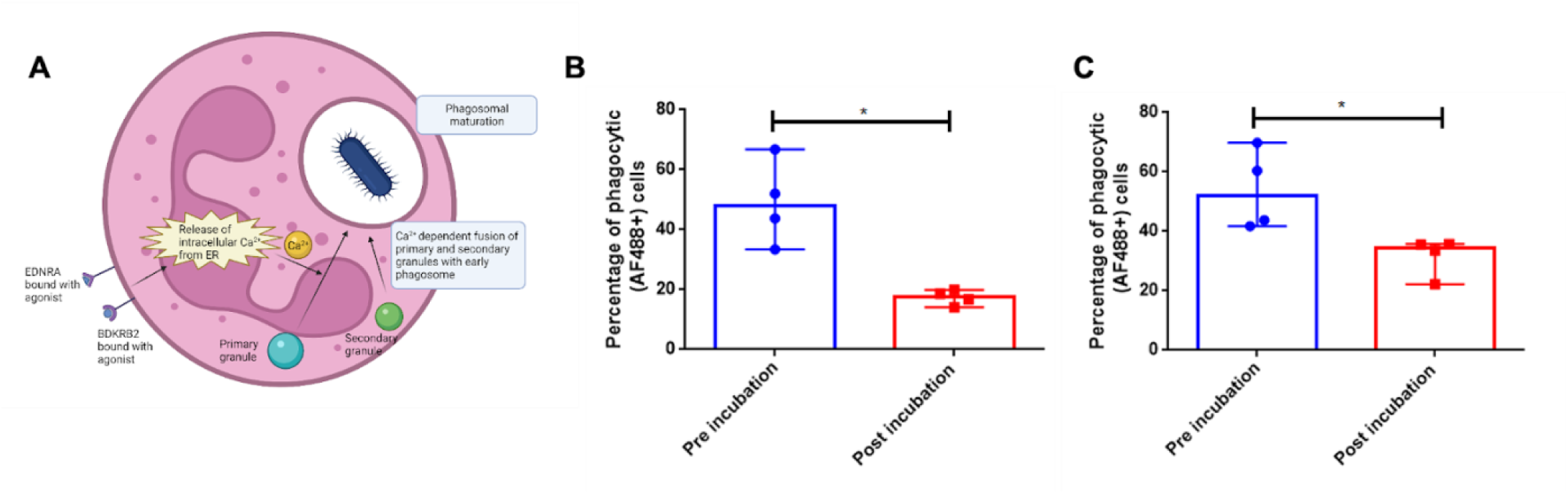
Effect of Alcohol on Phagocytosis. (A) Phagocytosis in neutrophils is dependent on calcium reserves. In neutrophils, elevated intracellular calcium levels facilitate the fusion of the primary and secondary granules to the early phagosome, thereby helping in phagosome maturation. (B) Graphical representation of percentage of phagocytic cells in pre-incubation vs alcohol incubation healthy samples (acute model). Median (Inter-Quartile Range) for pre-incubation sample group is 47.9 (36.0-63.1)%, and alcohol incubation sample group is 17.7 (14.78-19.52)%. Mann Whitney test was applied to compute the difference between the two groups [p value= 0.03] (n= 4 per group) (C) Graphical representation of percentage of phagocytic cells in pre-incubation vs alcohol incubation CLD samples (acute-on-chronic model). Median (Inter-Quartile Range) for pre-incubation sample group is 42.3 (41.8-52.10)%, and alcohol incubation sample group is 34.55 (25.05-35.73)%. Mann Whitney test was applied to compute the difference between the two groups. [p value= 0.03] (n= 4 per group).

### 3.4 IN SILICO MOLECULAR DOCKING OF ALCOHOL WITH CCL20 AND IL1β

Analysis of the nature of interactions between ethanol and *CCL20*, *IL1β* revealed three types of bonds which are primarily involved in the receptor-ligand binding: conventional hydrogen bonds, hydrophobic interactions and Van der Waals interactions. Binding energies of the ligand with receptors were less than −2.95 Kcal/mole and −2.85 Kcal/mole for *IL1β* and *CCL20* respectively, with the negative values indicating spontaneous binding and a higher negative value indicating increased binding affinity between the ligand and the receptor. The results of molecular docking have been summarized and both the 2D and 3D images of the docked complexes have been shown in Figure 7 (A-E).

**Figure 7.**
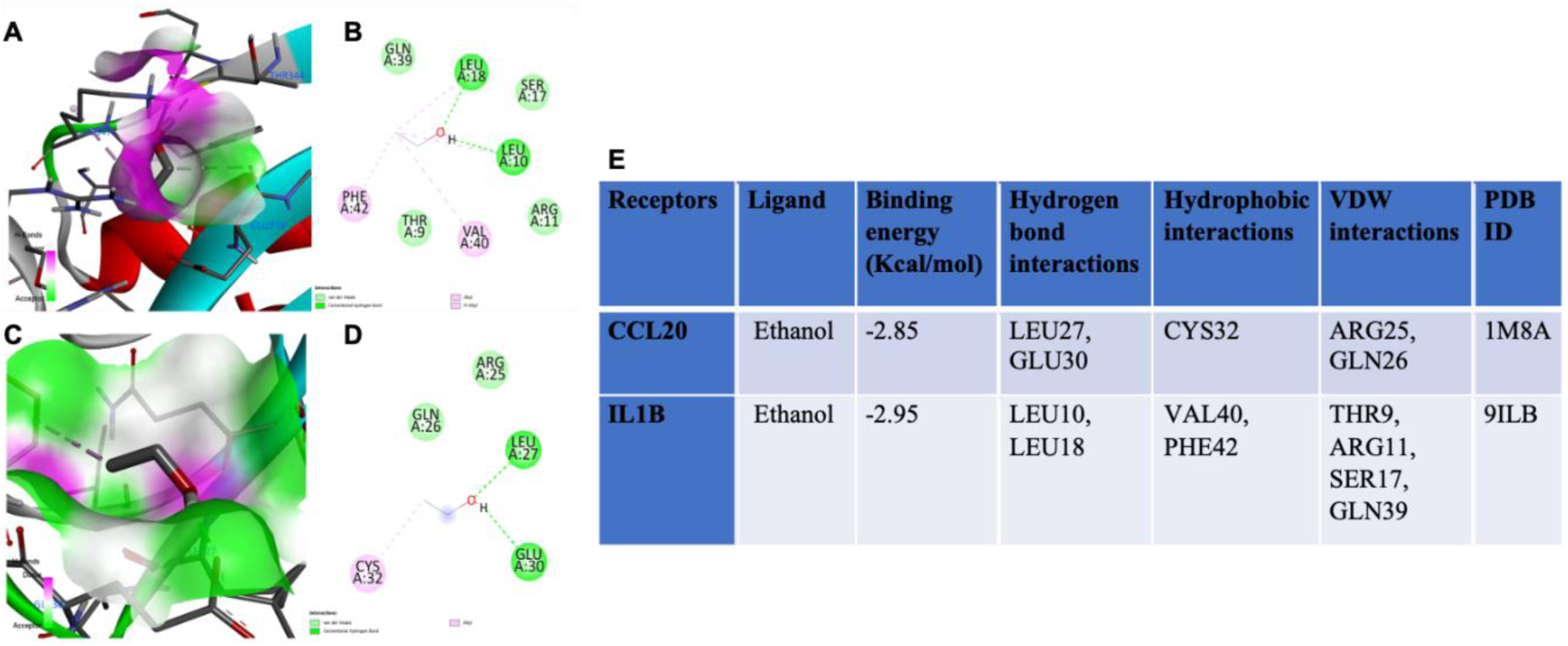
Three dimensional and two-dimensional representations of IL1β-Ethanol and CCL20-Ethanol docked complexes along with binding energy table. (A) Representation of the 3D and, (B) 2D docked complexes of IL1β with Ethanol as the ligand. (C) Representation of the 3D and, (D) 2D docked complexes of CCL20 bound with Ethanol. (E) Table showing the binding energies in KCal/mol of ethanol with CCL20 and IL-1β, the nature of interactions involved and the corresponding PDB IDs of the receptor proteins. All the dockings were performed using Autodock 4.2 software and the visualizations of the docked complexes were carried out using BIOVIA Discovery Studio Visualizer v21.1.0.20298. Dashed lines in green represent conventional hydrogen bonds whereas those in magenta represent hydrophobic interactions. Residues shown in faded green represent those involved in Van der Waals interaction with the ligand. Each docked complex has been supplemented with a receptor groove showing the binding site of ethanol where the green and pink shaded regions indicate localization of hydrogen bond acceptors and donors, respectively.

### 3.5 OVERLAPPING GENE EXPRESSION PATTERNS ACROSS CIRCULATING NEUTROPHILS IN ACLF AND PUBLISHED LIVER TISSUE TRANSCRIPTOME OF ALCOHOLIC LIVER DISEASE

In order to identify any overlapping alcohol responsive genes which are relevant during alcohol induced liver injury, a comparative analysis of our neutrophils (Alcoholic ACLF vs Non-Alcoholic ACLF) transcriptome dataset containing 283 DEG was performed with published Alcoholic liver datasets GSE28619 (34 genes), GSE155907 (4063 genes) and DisGeNET (195 genes) (Supplementary Figure S3-S4). The common DEG in alcoholic ACLF patient derived neutrophils and the liver tissue derived from patients of alcoholic liver disease (published transcriptomes) were: CCL20, IL1β, BDKRB2, ITKPA2, EDNRA1, TACR, among many others (Figure 8 A, B) (https://bioinfogp.cnb.csic.es/tools/venny/). Figure 8C shows shared genes between the alcoholic neutrophils DEG with the DisGeNET database. This database contains the disease-gene association from all curated, data mined, experimental and other published sources. The search keyword was: “Alcoholic liver disease” and the resulting gene set was downloaded. The overlapped DEG between Alcoholic neutrophils (our previously published) versus DisGeNET were a count of 5 genes (1.1%).

**Figure 8.**
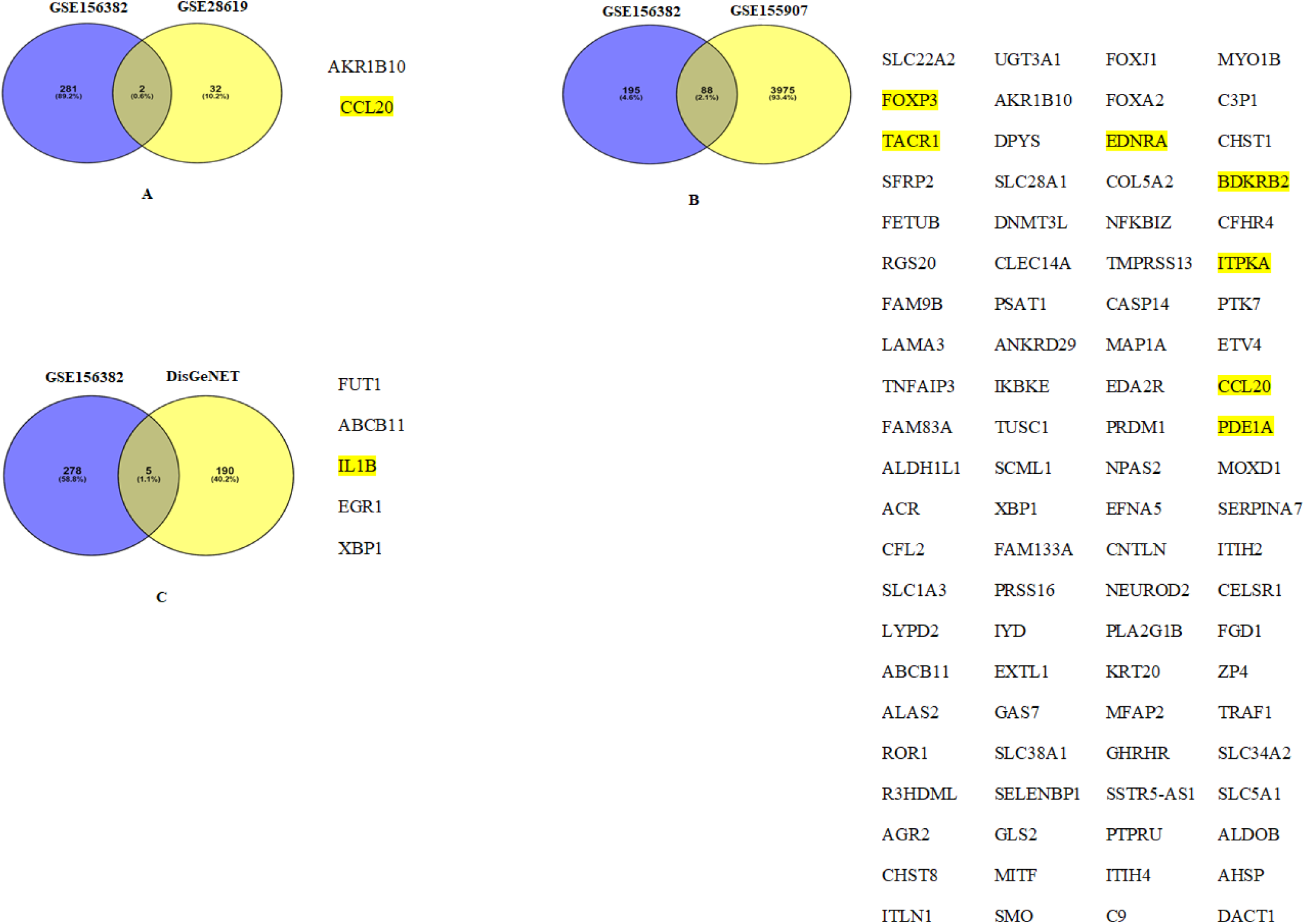
Representation of overlapping genes between Alcoholic ACLF PMN transcriptomics (GSE 156382 experimental dataset) and literature curated liver tissue transcriptome datasets. (A) GSE 156382 vs GSE 28619. This comparison resulted in 2 overlapping genes. (B) GSE 156382 vs GSE 155907. This comparison resulted in 88 overlapping genes. (C) GSE 156382 vs DisGeNet. This comparison resulted in 5 overlapping genes. Genes which were in common with our analysis included *CCL20, BDKRB2, ITPKA and IL1β* (highlighted in yellow).

## 4. DISCUSSION

Our analysis suggests that alcohol modulates gene expression and function of neutrophils. Alcohol is one of the most common etiologies of both chronic disease and acute injury in ACLF [10]. Analysis of previously published neutrophils transcriptome data revealed the overexpression of the inflammatory cytokine genes *CCL20* and *IL-1β* (Supplementary Table S1) and downregulation of the calcium signaling pathway (*BDKRB2, CCKAR, EDNRA, ERBB4, GRM5, ITPKA, PDE1A, TACR1*) in alcoholic-ACLF patient derived neutrophils (Figure 2). The calcium signalling pathway is central to the functional activation of neutrophils.[28]. Calcium signalling has been shown to be associated with downstream functions such as ROS generation and phagocytosis, both of which are anti-bacterial responses [28]. Therefore, a decrease in calcium signalling pathway might mean a corresponding decrease in ROS generation, phagocytosis and therefore an increased susceptibility to bacterial infection. The neutrophil transcriptome analysis in the alcoholic-ACLF vs non-alcoholic ACLF dataset suggested that the upregulation of inflammatory cytokine genes *CCL20* and *IL-1β* with the concomitant downregulation of calcium signalling pathway leading to defective ROS and phagocytosis might constitute the neutrophil dysfunction in alcoholic-ACLF.

Further, we optimized two different *in vitro* models to study the effects of alcohol on neutrophils-(i) an acute alcoholic model in which neutrophils derived from healthy donors were used and, (ii) an *in vitro* acute-on-chronic alcoholic model in which neutrophils enriched derived from chronic liver disease patients were used. These neutrophils were subsequently incubated with 300 mg/dL alcohol. As reported in published literature, blood alcohol levels of 300 mg/dL and above reflects severe alcohol intoxication and hence this dosage was selected in our *in vitro* model [30]. We carried out qRT-PCR analysis of two highly upregulated genes identified in alcoholic-ACLF patients from the preliminary microarray analysis-*IL-1β* (Log2 Fold = 1.65) and *CCL20* (Log2fold = 3.29) in the alcoholic-neutrophils model (Figure 2A, Supplementary Table S1). *IL-1β* and *CCL20* genes were also represented in the gene ontology analyses from published liver transcriptome analyses in alcoholic liver disease as well as in the overlap analysis with our dataset (Supplementary Figure S3 and Figure 8 respectively). In addition, *IL1β* is known to be a potent inflammatory cytokine that is produced as a result of activation of innate immune cells which leads to the activation of the NLRP3 inflammasome and production of *IL1β* [30]. Hence, these two genes were selected as candidates for further investigation as alcohol responsive genes. In our study, both *IL1β* and *CCL20* genes were significantly upregulated upon alcohol treatment of healthy neutrophils in the acute alcohol model (Figure 3 A-B), whereas only CCL20 upregulation was observed in alcohol treated CLD neutrophils in the acute-on-chronic alcohol model (Figure 3 C-D). We found that while the alcoholic neutrophils model showed elevation of either or both *IL1β* and *CCL20* genes upon treatment with alcohol, there was no difference between the gene expression values of alcoholic ACLF-derived neutrophils vs non-alcoholic ACLF derived neutrophils in the qRT-PCR cohort (Figure 3 G-H). This can be explained by the fact that both *IL1β* and *CCL20* are genes whose expression is regulated by multiple transcription factors and enhancer binding elements (Figure 3 I-J). Thus, alcohol might be one of the factors associated with IL1*β* and *CCL20* upregulation, but it is evident from our *in vitro* model that healthy and CLD neutrophils behave differently to alcohol treatment in terms of gene expression of IL1*β*.

Further, when we looked at neutrophil phenotypic changes, we observed that there was a distinct difference in healthy and CLD neutrophil ROS activation, both at baseline and stimulated, upon treatment with alcohol. We found that the healthy neutrophils at baseline did not generate significant ROS but PMA stimulation activated more than 90% of these neutrophils to generate ROS. Upon treatment with alcohol, healthy neutrophils generated significantly lower ROS in spite of PMA stimulation (Figure 5 B-C, Table 2). On the contrary, CLD neutrophils behaved similar to healthy at baseline, but had a major defect in the ability to cause activation when PMA was added. Further, alcohol had no significant effect on ROS generation in CLD neutrophils (Figure 5 D-E, Table 2). Therefore, our study showed that CLD neutrophils were more resistant to ROS production than healthy neutrophils. We also found that both acute and acute-on-chronic alcoholic neutrophils demonstrated decreased phagocytosis as assessed by flow cytometry (Figure 6, Table 2). Decreased neutrophils phagocytosis has been demonstrated in ACLF patient derived neutrophils in previously published studies [31,32].

**Table 2.** Neutrophil phenotype assays in response to alcohol using an *in vitro acute and acute-on-chronic* alcoholic neutrophil model. A-B: Acute alcohol model (n=4); C-D: Acute-on-chronic alcohol model (n=4). (A, C) Flow cytometry-based estimation of Reactive oxygen species (ROS), Phagocytosis and Intracellular Calcium Reserves. ROS estimations are represented as percentage (%) of ROS generating cells and also Mean fluorescent intensity (MFI). Phagocytosis is estimated as the percentage of total neutrophil cells which have engulfed the labeled *E. coli* bioparticles and generate a positive AF488 signal. Intracellular calcium reserves are measured as the difference in Fluo-4 dye (FITC) MFI from 0 minutes to 30 minutes. (B, D) Mann-Whitney test was performed to calculate the p-value significance for each assay, using GraphPad Prism software. P-value * indicates < 0.05.

| Sample ID | Pre-Incubation |  |  |  | Post-Incubation |  |  |  |
| --- | --- | --- | --- | --- | --- | --- | --- | --- |
|  | ROS % cells (MFI) |  | % Phagocytic cells | Calcium Reserve | ROS % cells (MFI) |  | % Phagocytic cells | Calcium Reserve |
|  | Only DHR/ DHR+PMA |  |  |  | Only DHR/ DHR+PMA |  |  |  |
| 1. | 7.41%<br>(3386) | 99.70%<br>(32571) | 52.00% | 53287 | 19.50%<br>(1161) | 62.50% (4521) | 14.10% | 22793 |
| 2. | 0.80%<br>(1451) | 99.60%<br>(37529) | 43.80% | 62832 | 36.50%<br>(1804) | 78.80% (2934) | 18.60% | 21967 |
| 3. | 5.07%<br>(2076) | 100% (44440) | 66.80% | 54853 | 1.84% (1342) | 25.30%<br>(14686) | 16.80% | 9558 |
| 4. | 13.5%<br>(3070) | 99.6%<br>(35850) | 33.40% | 41443 | 14.8% (1218) | 30.2% (3739) | 19.90% | 17265 |

B.
| Assays | P value |
| --- | --- |
| Baseline ROS cell % (only DHR 123) | 0.20 |
| Baseline ROS MFI (only DHR 123) | 0.06 |
| Stimulated ROS cell % (with PMA) | 0.03 |
| Stimulated ROS MFI (with PMA) | 0.03 |
| Calcium Reserve (MFI) | 0.03 |
| Phagocytosis cell % | 0.03 |

C.
| Sample ID | Pre-Incubation |  |  |  | Post-Incubation |  |  |  |
| --- | --- | --- | --- | --- | --- | --- | --- | --- |
|  | ROS % cells (MFI) |  | % Phagocytic cells | Calcium Reserve | ROS % cells (MFI) |  | % Phagocytic cells | Calcium Reserve |
|  | Only DHR/ DHR+PMA |  |  |  | Only DHR/ DHR+PMA |  |  |  |
| 1. | 11.20%<br>(1764) | 35.20%<br>(2529) | 41.80% | 42679 | 7.09% (1884) | 60.90%<br>(3902) | 35.80% | 9817 |
| 2. | 7.97% (1728) | 68.40%<br>(3746) | 43.80% | 76179 | 29.80%<br>(1842) | 15.40%<br>(4082) | 33.60% | 11973 |
| 3. | 3.76% (1606) | 59.80%<br>(2697) | 60.40% | 41345 | 7.02% (1544) | 64.90%<br>(1811) | 35.50% | 14077 |
| 4. | 6.04% (2095) | 47.80%<br>(3785) | 69.80% | 60608 | 15.40%<br>(1541) | 53.90%<br>(4133) | 22.20% | 12144 |

D.
| Assays | P value |
| --- | --- |
| Baseline ROS cell % (only DHR 123) | 0.34 |
| Baseline ROS MFI (only DHR 123) | 0.66 |
| Stimulated ROS cell % (with PMA) | >0.99 |
| Stimulated ROS MFI (with PMA) | 0.34 |
| Calcium Reserve (MFI) | 0.03(*) |
| Phagocytosis cell % | 0.03(*) |

In neutrophils, both ROS generation and phagocytosis are downstream to calcium signalling. The transcriptome of alcoholic-ACLF patients indicated a downregulation of the calcium signalling pathway (Figure 2D) therefore, we assessed the calcium reserve in alcoholic-neutrophils and compared them with baseline healthy and CLD neutrophils. The calcium assay in this study measures intracellular concentrations of Ca^2+^ over time. Our results showed that both healthy and CLD alcohol treated neutrophils had lower calcium concentrations post alcohol treatment suggesting that alcohol treatment reduced intracellular Ca^2+^ reserves (Figure 4, Table 2). This may lead to the dampening of calcium signalling pathway which can influence both phagocytosis and ROS generation thereby leading to defective anti-bacterial functions and increased susceptibility to infection [31]. This increased susceptibility to bacterial infections is reported in ACLF as well as alcoholic liver disease[33,35]. While the importance of this pathway in regulation of neutrophils as well as monocyte functions is well established, the molecular cues that regulate calcium signalling in these innate immune cells are not understood. Our study suggests that alcohol may be one of the molecular cues that downregulates calcium signalling in innate immune cells.

Alcohol is known to form protein adducts through its metabolic aldehyde intermediate causing cellular cytotoxicity therefore, we wanted to examine if it can also directly bind any of the upregulated inflammatory proteins [36]. Therefore, we carried out *in silico* molecular docking studies of ethanol with PDB structures of *CCL20* and *IL1β* (Figure 7 A-D). Molecular docking involving proteins and ligands evaluate their potential to interact as well as reveal the nature of interactions, besides serving as a tool for visualization of these interactions [37]. Our analysis revealed strong ethanol binding with *IL1β* with a binding energy of −2.95 KCal/mol and with *CCL20* with a binding energy of −2.85 Kcal/mol, with the involvement of hydrogen bonds, hydrophobic interactions and Van der Waals interactions. In molecular docking analyses, negative binding energy indicates spontaneous interaction and therefore this analysis suggests that in addition to its cellular effects on neutrophils, alcohol may also directly interact with the *CCL20* and *IL1β* proteins. This is a novel observation warranting detailed biophysical measurements which can be explored in the future.

Re-analysis of published liver tissue transcriptome datasets from patients of alcoholic liver disease was carried out in order to investigate the relevance of the neutrophil genes identified in our study. The datasets GSE28619, GSE155907 (published liver tissue transcriptomic datasets) and GSE156382 (our neutrophil transcriptome dataset) shared the upregulation of *CCL20* and *IL1β* genes in patients with alcoholic liver disease (Figure 8). *IL1β* is a key mediator of inflammation and ACLF patients have been shown to have elevated plasma levels of the cytokine. *IL1β* is known to be a master regulator of inflammation and induces acute phase responses [38]. These data therefore suggest that the increase in *CCL20* and *IL1β* which can occur due to the effect of alcohol on neutrophils, are also relevant in liver injury in alcoholic liver disease possibly by enhancing inflammatory cell damage within the liver tissue possibly in response to alcohol.

## 5. CONCLUSIONS

Our study showed that the expression of the genes *CCL20* and *IL1β* increased upon alcohol exposure of healthy neutrophils with concomitant dampening of calcium signaling, ROS production, and phagocytosis. Our data also showed that neutrophils derived from chronic liver disease (CLD) patients behaved differently from those derived from healthy patients in terms of gene expression as well as functional assays. This suggested that there are underlying differences in chronic liver disease derived neutrophils and neutrophils from healthy individuals that warrant further investigations.

## Supporting information

MiQE docx

## Supplementary Materials

Figure S1: PMN enrichment measurement by flow cytometry (n=4; representative); Figure S2: Viability assay of enriched PMN using Zombie Aqua fixable dye; Figure S3: WebGestalt based analysis of Liver tissue transcriptomic datasets GSE 28619 and GSE 155907; Figure S4: Curation of ALD Tissue Transcriptomic Data; Table S1: List of DEG’s Alcoholic v/s Non-Alcoholic Neutrophils; Table S2: Pathway analysis of differentially expressed genes in neutrophils isolated from ACLF patients with alcohol etiology; Table S3: Baseline characteristics of ACLF (n=30) patients, CLD (n=10) and HC (n=15) patients used in validation experiments (qRT-PCR).

## Author Contributions

The authors confirm contribution to the paper as follows. Study conception and design: PA, RS, DC, TL. Data collection: RS, DC, TL, AR, SS. Analysis and interpretation of results: PA, RS, DC, TL, AR, SS, S. Draft manuscript preparation: RS, DC, PA, TL, AR, SS, S.

## Informed Consent Statement

The studies involving human participants were reviewed and approved by 1. All India Institute of Medical Sciences, New Delhi ethics committee [Reference No. IEC/473/9/2016 and, IEC/369/7/2016], 2. All India Institute of Medical Sciences, New Delhi ethics committee the Institute Ethics Committee (Ref. No.IEC/687/8/2019). The patients/participants provided their written informed consent to participate in this study.

## Data Availability Statement

The original contributions presented in the study are included in the article/supplementary materials. The data presented in this study are available in [7]. Further inquiries can be directed to the corresponding author.

