## Supplementary Figure S1 for "Alcohol Responsive Genes in the Neutrophils of Acute-On-Chronic Liver Failure Reveal Modulation of Intracellular Calcium, ROS, and Phagocytosis"

### Slide 1
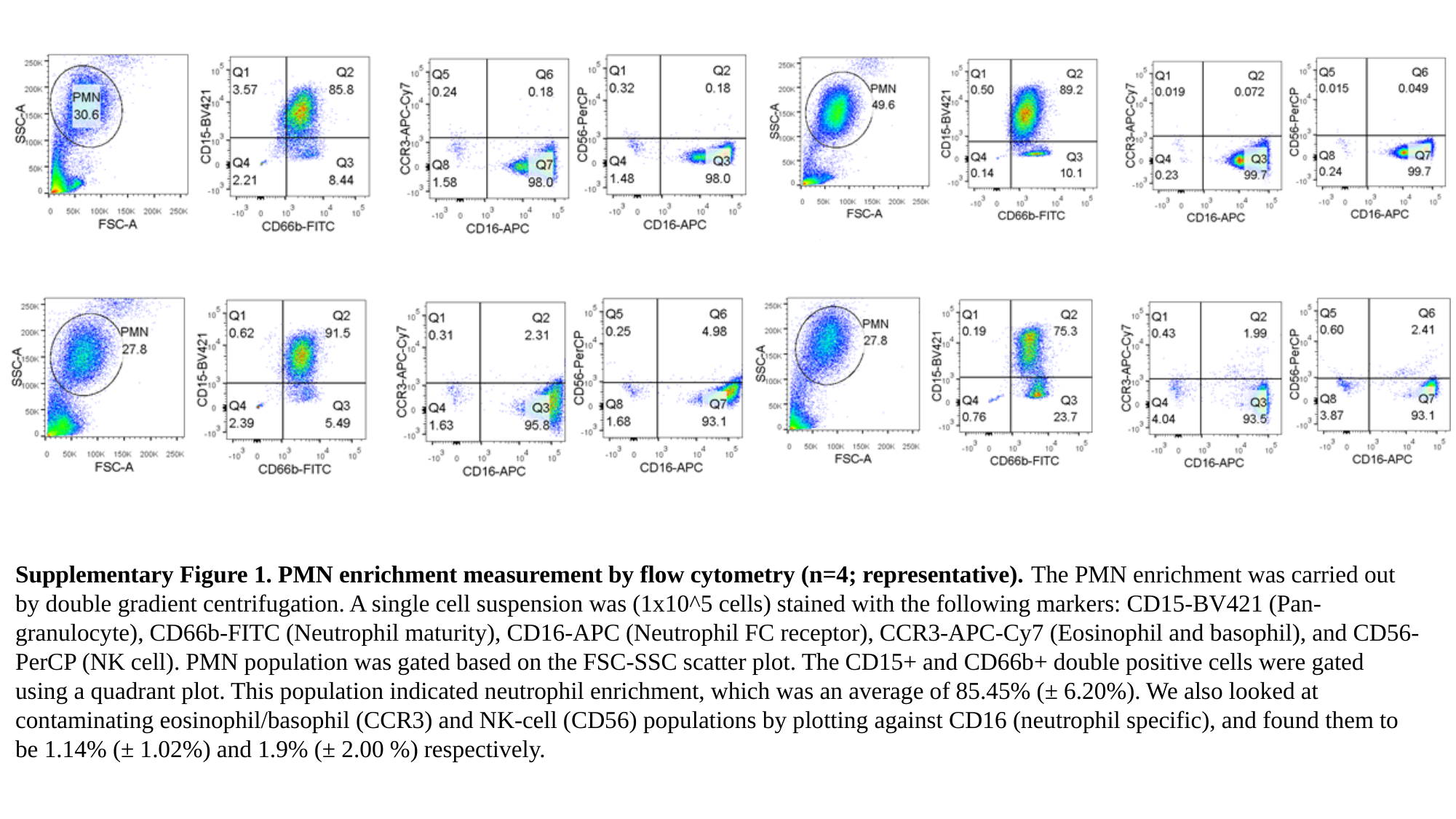

Supplementary Figure 1. PMN enrichment measurement by flow cytometry (n=4; representative). The PMN enrichment was carried out by double gradient centrifugation. A single cell suspension was (1x10^5 cells) stained with the following markers: CD15-BV421 (Pan-granulocyte), CD66b-FITC (Neutrophil maturity), CD16-APC (Neutrophil FC receptor), CCR3-APC-Cy7 (Eosinophil and basophil), and CD56-PerCP (NK cell). PMN population was gated based on the FSC-SSC scatter plot. The CD15+ and CD66b+ double positive cells were gated using a quadrant plot. This population indicated neutrophil enrichment, which was an average of 85.45% (± 6.20%). We also looked at contaminating eosinophil/basophil (CCR3) and NK-cell (CD56) populations by plotting against CD16 (neutrophil specific), and found them to be 1.14% (± 1.02%) and 1.9% (± 2.00 %) respectively.
