## Supplementary Figure S2 for "Alcohol Responsive Genes in the Neutrophils of Acute-On-Chronic Liver Failure Reveal Modulation of Intracellular Calcium, ROS, and Phagocytosis"

### Slide 1
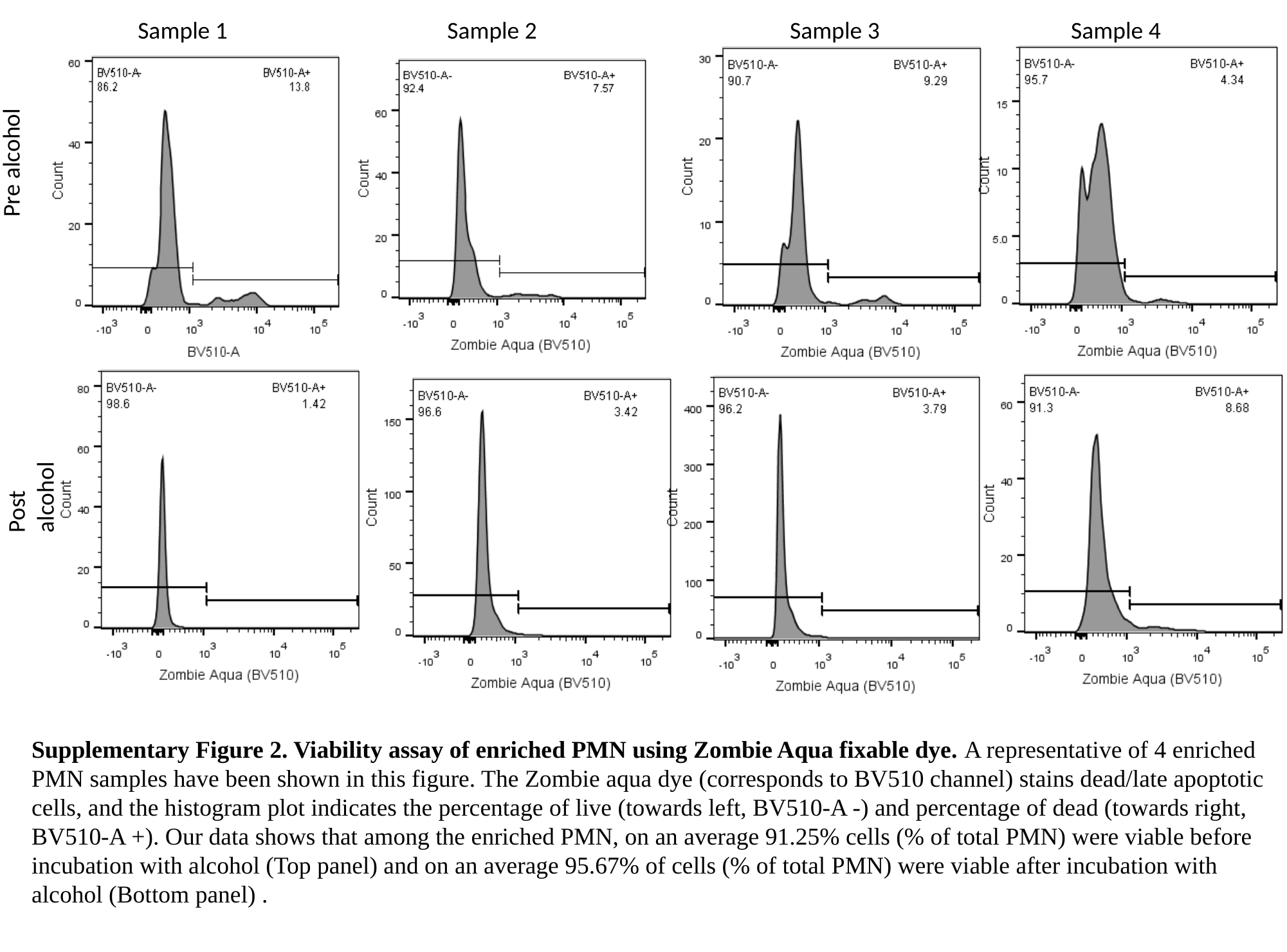

Sample 1
Sample 2
Sample 3
Sample 4
Pre alcohol
Post alcohol
Supplementary Figure 2. Viability assay of enriched PMN using Zombie Aqua fixable dye. A representative of 4 enriched PMN samples have been shown in this figure. The Zombie aqua dye (corresponds to BV510 channel) stains dead/late apoptotic cells, and the histogram plot indicates the percentage of live (towards left, BV510-A -) and percentage of dead (towards right, BV510-A +). Our data shows that among the enriched PMN, on an average 91.25% cells (% of total PMN) were viable before incubation with alcohol (Top panel) and on an average 95.67% of cells (% of total PMN) were viable after incubation with alcohol (Bottom panel) .
