## Supplementary Figure S3 for "Alcohol Responsive Genes in the Neutrophils of Acute-On-Chronic Liver Failure Reveal Modulation of Intracellular Calcium, ROS, and Phagocytosis"

**Supplementary Figure 3. Webgestalt based analysis of Liver tissue transcriptomic datasets GSE 28619 and GSE 155907.** Gene ontology analysis was performed for upregulated and downregulated genes separately. Gene set enrichment analysis (GSEA) was performed using the entire differentially expressed gene list. (A) Upregulated genes for GSE 28619 dataset. (B) Downregulated genes for GSE 28619 dataset. (C) GSEA for GSE 28619 dataset. (D) Upregulated genes for GSE 155907 dataset. (E) Downregulated genes for GSE 155907 dataset. (F) GSEA for GSE 155907 dataset.

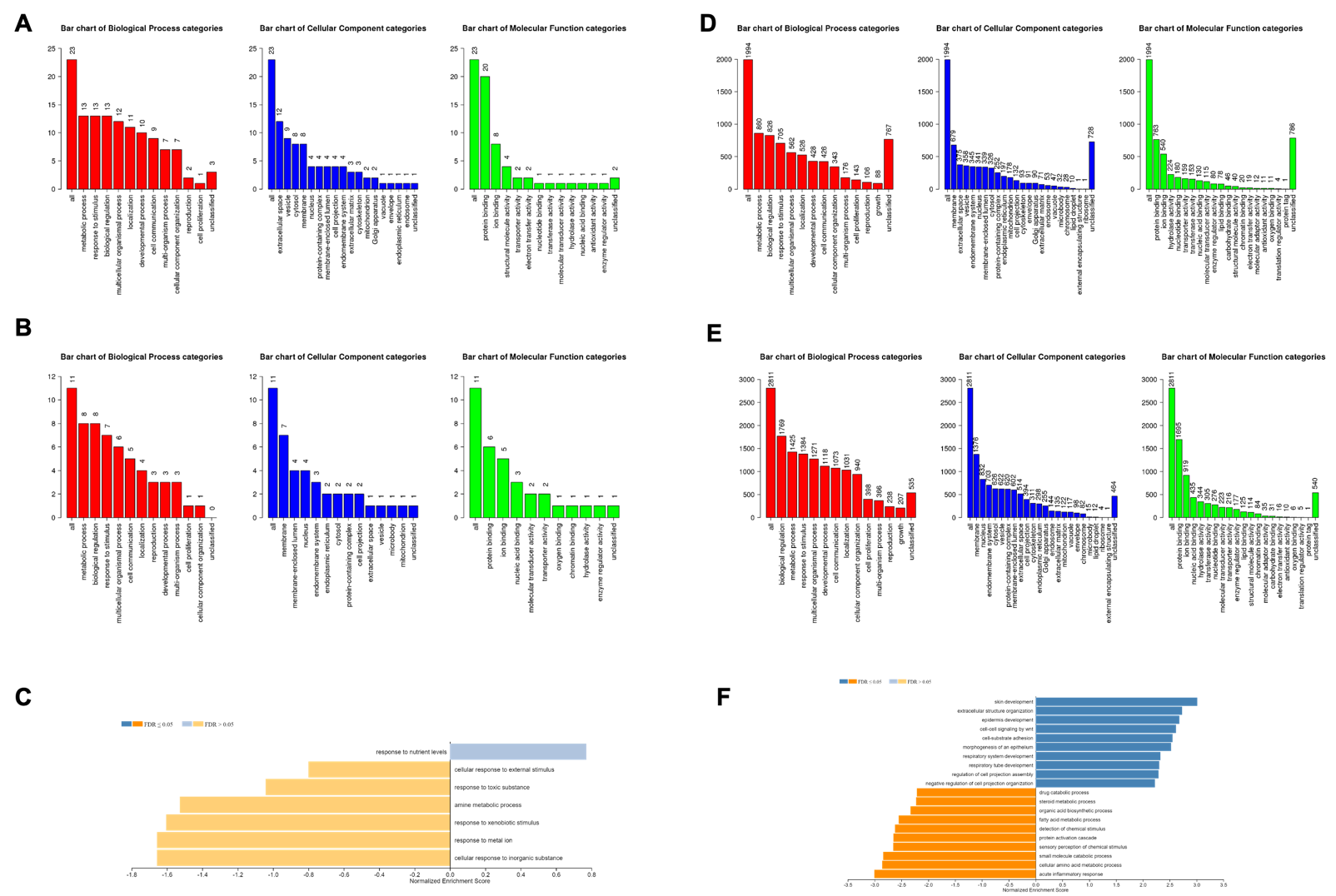
