## Supplementary figures and images for "Alcohol Responsive Genes in the Neutrophils of Acute-On-Chronic Liver Failure Reveal Modulation of Intracellular Calcium, ROS, and Phagocytosis"

### Supplementary Figure S4

#### Slide 1
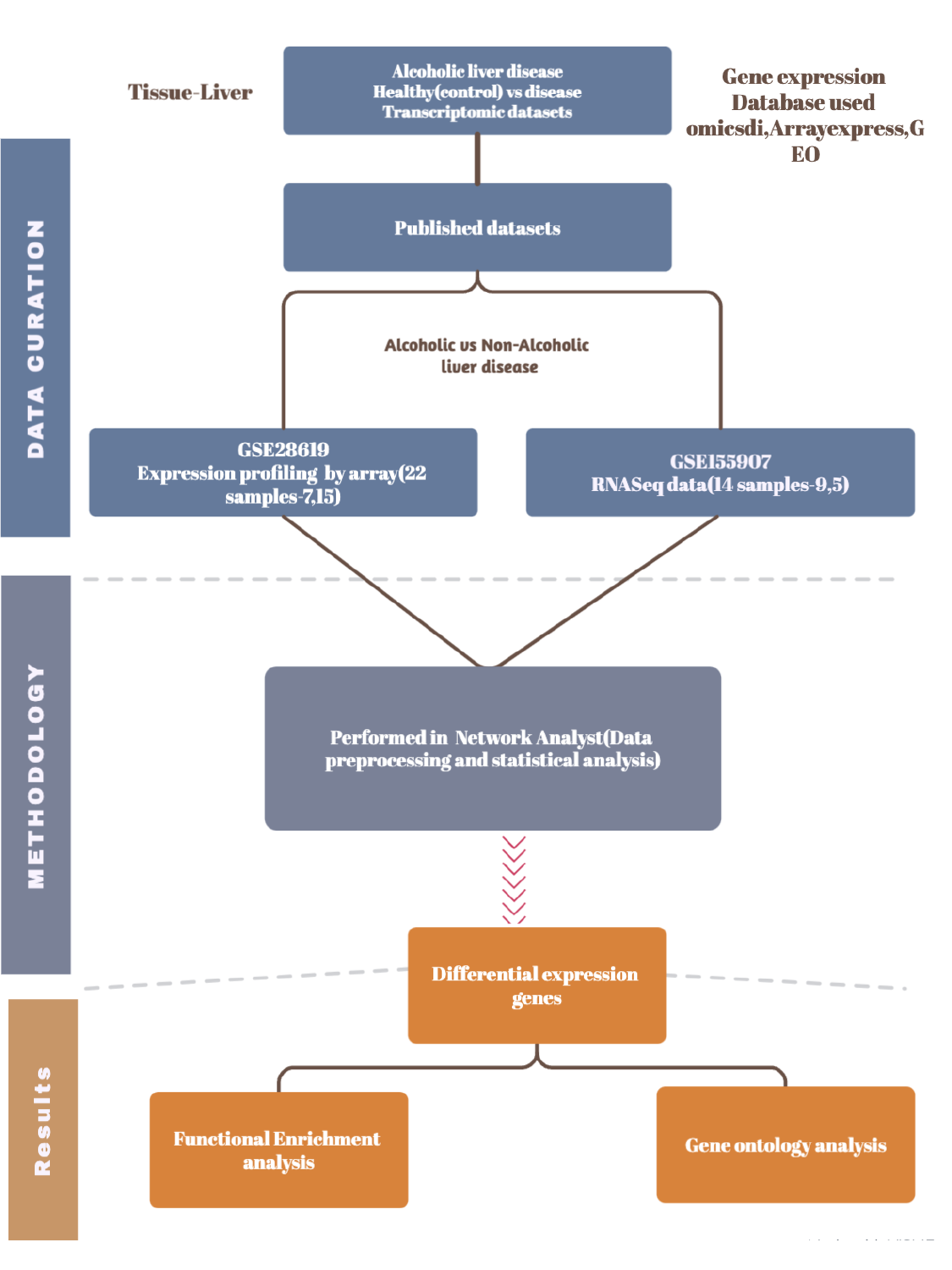
