## Supplementary Table S2 for "Alcohol Responsive Genes in the Neutrophils of Acute-On-Chronic Liver Failure Reveal Modulation of Intracellular Calcium, ROS, and Phagocytosis"

**Table 2**. **Pathway analysis of differentially expressed genes in neutrophils isolated from ACLF patients with alcohol etiology.**

Differentially expressed genes (DEG) from neutrophils from alcoholic ACLF patients were subjected to pathway enrichment analysis using Metascape tool. The table above shows the most significantly enriched pathways, their corresponding IDs, logP values and the list of genes upregulated and downregulated in each pathway.

| Pathway | ID | Number of DEG involved | Log p-value | Upregulated genes | Downregulated genes |
| --- | --- | --- | --- | --- | --- |
| T-cell differentiation | GO:0030217 | 42 | -6.47022 | PRDM1, EGR1  , EGR3, GPR18, IL1B, PTPN2, XBP1, NFKBIZ, LOXL3, SLAMF6, CX3CR1, APPL1, TNFAIP3, SLAMF7,  IKBKE | FOXJ1, PLA2G2D, FOXP3, IL21,  EFNA5, FUT1, FOXA2, KDR, LAMA3, MELTF, SFRP2, PTPRU, AGR2, CSPG5, ZP4, EGFLAM, APOA4, MITF, SLC22A13, C9, PGLYRP3, CRLF2, HOXA5, PLA2G1B, CDH3, FLT4, ACKR1 |
| GPCR Signalling | R-HSA-500792 | 27 | -5.26844 | CX3CR1, GPR18, CCL20,  GPR84 | BDKRB2, CCKAR, EDNRA, FSHR, ACKR1, GHRHR, MLNR, GRM3, GRM5, PPY, RHO, SMO, TACR1, UTS2, FGD1, PDE1A, RGS20, ERBB4, ITPKA, OR2W1, CASQ1, PLA2G1B, PRKD1 |
| Calcium Signalling | hsa04020 | 8 | -3.2926 |  | BDKRB2, CCKAR, EDNRA, ERBB4, GRM5, ITPKA, PDE1A, TACR1 |
| Neurotransmitter release | R-HSA-112310 | 27 | -2.7056 | SLC1A3, KCNJ2, SLC38A1, IL1B, PSAT1, SLCO4A1 | CHAT, SLC18A2, SLC22A2, PPFIA3, GLS2, ERBB4, GRM5, KCNA7, KCNJ9, KCNQ4, SLITRK2, CSPG5, MOXD1, SLC5A1, SLC28A1, SLC34A2, BDKRB2, EDNRA, SNTA1, UTS2, GRM3 |
| Transmembrane RTK | GO:0007169 | 38 | -5.21547 | IL1B, PTPN2, APPL1, EGR1, TNFAIP3, CX3CR1, PRDM1, EGR3 | BDKRB2, CDH3, EFNA5, ERBB4, FGF4, FLT4, GHRHR, IGFBP4, KDR, ROR1, PRKD1, AGR2, DNAI1, ANGPT4, FAM83A, CLEC14A, SHC4, DUSP7,  GRM5, FOXA2, PFKFB1, PLA2G1B, SFRP2, ITLN1, IL21, CLDN19, PLA2G2D, COL5A2, LAMA3, PTPRU |
| Cytokine signalling | GO:0019221 | 37 | -4.04544 | CX3CR1, EGR1, IL1B, PTPN2, CCL20, TNFAIP3, TRAF1, IKBKE,  APPL1, EGR3,  GPR18 | ACKR1, CCL26, IL36A, EDA2R, CRLF2, DCST1, DUSP7, ERBB4, FGF4, FLT4, FSHR, IGFBP4, KDR, MOS, ROR1, PLA2G1B  SFRP2, DACT1, EFNA5, ETV1,  LAMA3, PRKD1, RHO, SMO, ROBO4, IL21 |
