## Supplementary Table S3 for "Alcohol Responsive Genes in the Neutrophils of Acute-On-Chronic Liver Failure Reveal Modulation of Intracellular Calcium, ROS, and Phagocytosis"

|  | **ACLF (n=30)** | **CLD (n=10)** | **HC (n=15)** | **P value** |
| --- | --- | --- | --- | --- |
| **Age, years, Mean ± SD** | 48 ± 13 | 40 ± 9 | 39 ± 12 | 0.78 |
| **Males %** | 70% | 80% | 66% | 0.0758 |
| **Hemoglobin (g/dL)** | 8.30 (7.50-8.80) | 11.20 (10.00-13.78) | n/a | 0.014 |
| **Sodium (mEq/L)** | 139.8 ± 2.09 | 137.8 ± 1.02 | n/a | 0.61 |
| **Albumin (g/dL)** | 2.65 ± 0.13 | 3.71 ± 0.45 | n/a | 0.0055 |
| **Total Leukocyte Count (x103 per mm3)** | 14.20 ± 1.92 | 6.77 ±0.56 | n/a | 0.04 |
| **Platelet (x103 per mm3)** | 80.84 ± 13.17 | 120.2 ± 11.95 | n/a | 0.12 |
| **Urea (mg/dL)** | 111.5 ± 22.29 | 22.25 ± 3.65 | n/a | 0.04 |
| **Creatinine (mg/dL)** | 1.60 (0.90-3.40) | 0.95 (0.65-1.20) | n/a | 0.016 |
| **Potassium (mEq/L)** | 4.55 ± 0.27 | 4.35 ± 0.35 | n/a | 0.70 |
| **Bilirubin (mg/dL)** | 10.69 ± 2.26 | 1.968 ± 0.53 | n/a | 0.04 |
| **Aspartate Aminotransferase (IU/L)** | 120 ± 24.92 | 81.17 ± 16.50 | n/a | 0.39 |
| **Alanine Transaminase (IU/L)** | 52.17 ± 11.02 | 75.33 ± 24.85 | n/a | 0.36 |
| **Serum Alkaline Phosphatase (IU/L)** | 293.90 ± 45.82 | 232.80 ± 35.56 | n/a | 0.47 |

|  | **Non-Alcoholic ACLF (n=15)** | **Alcoholic ACLF (n=15)** | **P value** |
| --- | --- | --- | --- |
| **Age, years, Mean ± SD** | 54 ± 4 | 42 ± 5 | 0.14 |
| **Males %** | 42.85% | 100% | < 0.0001 |
| **Hemoglobin (g/dL)** | 8.30 (7.50-8.30) | 8.60 (7.02-10.3) | 0.28 |
| **Sodium (mEq/L)** | 137.7 ± 3.29 | 141.7 ± 2.60 | 0.35 |
| **Albumin (g/dL)** | 2.69 ± 0.14 | 2.62 ± 0.24 | 0.85 |
| **Total Leukocyte Count (x103 per mm3)** | 10.73 ± 2.93 | 9.74 ± 3.62 | 0.83 |
| **Platelet (x103 per mm3)** | 88.89 ± 20.23 | 73.60 ± 17.89 | 0.57 |
| **Urea (mg/dL)** | 93.60 ± 29.48 | 131.40 ± 34.34 | 0.40 |
| **Creatinine (mg/dL)** | 2.40 (1.75-4.12) | 1.40 (0.57-1.65) | 0.02 |
| **Potassium (mEq/L)** | 4.80 ± 0.34 | 4.3 ± 0.40 | 0.37 |
| **Bilirubin (mg/dL)** | 9.10 ± 3.17 | 12.12 ± 3.31 | 0.52 |
| **Aspartate Aminotransferase (IU/L)** | 132.80 ± 42.13 | 111.5 ± 25.96 | 0.60 |
| **Alanine Transaminase (IU/L)** | 59.44 ± 19.34 | 44.89 ± 11.36 | 0.52 |
| **Serum Alkaline Phosphatase (IU/L)** | 252.00 ± 63.94 | 335.90 ± 66.28 | 0.38 |

**Supplementary Table 3. Baseline characteristics of ACLF (n=30) patients, CLD (n=10) and HC (n=15) patients used in validation experiments (qRT-PCR).** The 30 ACLF samples used in gene expression validation experiment, were also stratified as alcoholic ACLF (n=15) and Non-alcoholic ACLF (n=15). Baseline patient characteristics were compared and p-value was computed for each variable.
