## Supplementary material for "Alcohol Responsive Genes in the Neutrophils of Acute-On-Chronic Liver Failure Reveal Modulation of Intracellular Calcium, ROS, and Phagocytosis": MiQE docx

**I. Experimental design:**

(i) Experimental groups:

Chronic liver disease patients/Healthy individuals treated with alcohol (n=14)

(ii) Control groups:

Untreated chronic liver disease patients/Healthy individuals (n=14)

**II. Sample:**

**Volume/mass of sample processed:** 6 ml Blood samples were acquired in each study groups in EDTA coated purple top vacutainers. Plasma was aseptically separated and stored at − 80 °C until further use.

**PMN Isolation:** PMN were isolated by modified Boyum's method of double gradient centrifugation. Blood pellet containing the buffy coat was diluted with 2× volume of sterile 1× PBS (*VWR, USA, 97062-730*), at room temperature (RT). Ficoll-Hisep (*Himedia, INDIA, LSM 1077*) was layered over Granulosep (*Himedia, INDIA, LS004*) in a 2:3 ratio to prepare a double gradient in a 15 ml centrifuge tube. Whole blood was carefully layered on top followed by centrifugation at 300×*g* for 30 min at RT without brakes. The following phases were formed in order (top to bottom) after centrifugation: Diluted plasma, PBMC, Hisep, PMN, Granulosep, RBC pellet. Diluted plasma was discarded; PBMC layer was separated and the lower enriched PMN layer was collected. PMN cells were resuspended in sterile 1 × PBS and washed twice by centrifugation at 500×*g* for 10 min. Contaminating RBCs were removed by incubating the washed pellet in 1 × RBC lysis solution for 10 min at 4 °C, followed by two 1 × PBS washes. The final cell pellet was reconstituted in 1.2 ml filtered RPMI-1640 cell culture media (*Himedia, INDIA, AL171A*). Enriched neutrophils were seeded on a 6 well plate at a density of 1 x 10^5^ cells. Complete RPMI media was supplemented with 300 mg/dL of culture-grade 100% Ethanol. The plates were incubated for 24 hours at 5% CO2, and 37°C. Post Incubation, the neutrophils were harvested by centrifugation of the collected media and washed twice with incomplete RPMI.

**III. qPCR target information**

**(i) Primer designing strategy:** The primers against the set of referred genes were designed using NCBI primer BLAST software. The design parameters settings were chosen to all primers span the exon-exon junctions and intron-exclusion criteria. Also, primer designs were allowed to amplify mRNA splice variants. The specificity of designed primers was verified by BLAST platform of NCBI.

**(ii) Manufacturer of oligonucleotides:** IDT

| **Gene Name** | **Primer Sequence (5’->3)** | **Length** | **Amplicon Size** |
| --- | --- | --- | --- |
| **18S** | |  |  |
| Forward | GTAACCCGTTGAACCCCATT | 20 | 151 |
| Reverse | CCATCCAATCGGTAGTAGCG | 20 |  |
| **Il1β** |  |  |  |
| Forward | CCACAGACCTTCCAGGAGAATG | 22 | 131 |
| Reverse | GTGCAGTTCAGTGATCGTACAGG | 23 |  |
| **CCL20** | | | |
| Forward | GGACATAGCCCAAGAACAGAAA | 22 | 99 |
| Reverse | GTCCAGTGAGGCACAAATTAGA | 22 |  |

**IV. qPCR protocol**:

**(i) Complete reaction conditions:** SYBR green 1 chemistry was used to determine the relative expression of the target genes. The cycling parameters are: Initial denaturation at 95°C for 3 minutes, followed by annealing, amplification and detection for 40 cycles (95°C 30’’, 60°C 20’’, 72°C 20’’) at end-point fluorescence, and melt-curve analysis at 60-95°C at 0.1 °C rise per second at continuous fluorescence to detect specific amplicons.

**(ii) Reaction volume and components:** Reaction volume of total 10 µl : Components were 8 µl mastermix consisting of 5 µl of SYBR Green buffer. Mastermix (Thermo Fisher Scientific Dynamo Flash: F415S), 0.2 µl of forward + 0.2 µl of reverse primers (0.4mM), 2.6 µl of nuclease-free water, and 2 µl of template cDNA.

**(v) Manufacture of qPCR instruments:** Agilent AriaMx Real-Time PCR system was used to carry out the qPCR experiments.

**Keywords and abbreviations**

MIQE: The Minimum Information for Publication of Quantitative Real-Time PCR Experiments qPCR: Quantitative real time PCR

CLD: Chronic Liver Disease

DC: Disease controls

**PCR standardization and primer efficiency:**

18S rRNA was used as the housekeeping gene. Primer validations and standardization of the reaction conditions were done by temperature gradient PCR in Himedia Prima-96™ Well Block system. Reaction conditions were optimized to be same for all the genes, and chosen for qPCR as follows: 95°C 3’, 95°C 30’’, 60°C 20’’, 72°C 20’’. Primer efficiency was measured for each primer pair by the standard curve method, using a 2-fold dilution series of the pooled cDNA template from each of the groups: CLD and Healthy Control samples, by qPCR.

**18S**

**IL1**$\boldsymbol{\beta}$

**CCL20**

| Log Concentration | Cq Value |
| --- | --- |
| 2.39794 | 19.29 |
| 2.09691 | 20.325 |
| 1.79588 | 21.405 |
| 1.49485 | 22.21 |
| 1.19382 | 23.315 |

| Log Concentration | Cq Value |
| --- | --- |
| 2.39794 | 27.875 |
| 2.09691 | 28.99 |
| 1.79588 | 29.6 |
| 1.49485 | 30.59 |
| 1.19382 | 31.825 |

| Log Concentration | Cq Value |
| --- | --- |
| 2.39794 | 29.71 |
| 2.09691 | 30.835 |
| 1.79588 | 31.565 |
| 1.49485 | 32.3 |
| 1.19382 | 32.97 |

|  | **18S** | **IL1**$\boldsymbol{\beta}$ | **CCL20** |
| --- | --- | --- | --- |
| **EFFICIENCY %** | 100.91% | 107.43% | 117.52% |
| **LOG EFFICIENCY** | 2.003936603 | 2.031135798 | 2.070105465 |

**References:**

**(In accordance with the MIQE^1^ guidelines):**<http://clinchem.aaccjnls.org/content/55/4/611>
